# Extensive long-range polycomb interactions and weak compartmentalization are hallmarks of human neuronal 3D genome

**DOI:** 10.1101/2023.08.04.551939

**Authors:** Ilya A. Pletenev, Maria Bazarevich, Diana R. Zagirova, Anna D. Kononkova, Alexander V. Cherkasov, Olga I. Efimova, Eugenia A. Tiukacheva, Kirill V. Morozov, Kirill A. Ulianov, Dmitriy Komkov, Anna V. Tvorogova, Vera E. Golimbet, Nikolay V. Kondratyev, Sergey V. Razin, Philipp Khaitovich, Sergey V. Ulianov, Ekaterina E. Khrameeva

**Affiliations:** Center for Molecular and Cellular Biology, Skolkovo Institute of Science and Technology, Moscow, 121205, Russia; A.A. Kharkevich Institute for Information Transmission Problems, Moscow, 127051, Russia; Center for Neurobiology and Brain Restoration, Skolkovo Institute of Science and Technology, Moscow, 121205, Russia; Department of Biological and Medical Physics, Moscow Institute of Physics and Technology, Moscow, 141700, Russia; Department of Molecular Biology, Faculty of Biology, M.V. Lomonosov Moscow State University, Moscow, 119991, Russia; CNRS UMR9018, Institut Gustave Roussy, Villejuif, 94805, France; Koltzov Institute of Developmental Biology, Russian Academy of Sciences, Moscow, 119334, Russia; Department of Cellular Genomics, Institute of Gene Biology, Russian Academy of Sciences, Moscow, 119334, Russia; Center for Precision Genome Editing and Genetic Technologies for Biomedicine, Institute of Gene Biology, Russian Academy of Sciences, Moscow, 119334, Russia; Laboratory of Clinical Genetics, Mental Health Research Center, Moscow, 115522, Russia

**Author notes:** To whom correspondence should be addressed.; Correspondence may also be addressed to Sergey Ulianov. Joint Authors (Ilya A. Pletenev, Maria Bazarevich, Diana R. Zagirova).

## Abstract

Chromatin architecture regulates gene expression and shapes cellular identity, particularly in neuronal cells. Specifically, polycomb group (PcG) proteins enable establishment and maintenance of neuronal cell type by reorganizing chromatin into repressive domains that limit the expression of fate-determining genes and sustain distinct gene expression patterns in neurons. Here, we map the 3D genome architecture in neuronal and non-neuronal cells isolated from the Wernicke’s area of four human brains and comprehensively analyze neuron-specific aspects of chromatin organization. We find that genome segregation into active and inactive compartments is greatly reduced in neurons compared to other brain cells. Furthermore, neuronal Hi-C maps reveal strong long-range interactions, forming a specific network of PcG-mediated contacts in neurons that is nearly absent in other brain cells. These interacting loci contain developmental transcription factors with repressed expression in neurons and other mature brain cells. But only in neurons, they are rich in bivalent promoters occupied by H3K4me3 histone modification together with H3K27me3, which points to a possible functional role of PcG contacts in neurons. Importantly, other layers of chromatin organization also exhibit a distinct structure in neurons, characterized by an increase in short-range interactions and a decrease in long-range ones.

## INTRODUCTION

Mammalian genomes possess a complex 3D architecture reflecting a composite interplay between structure and functionality. Structures of different scales appear to be formed by distinct mechanisms and presented hierarchically within each other (1). Traditionally, four levels of chromatin organization are distinguished: chromosome territories, compartments, Topologically Associating Domains (TADs), and chromatin loops. Chromosome territories are distinct nuclear areas, preferentially occupied by different interphase chromosomes. At coarse-grain level, chromosomes are segregated into two compartments, A and B, comprising regions of similar epigenetic states - “active” and “inactive” chromatin, respectively (2). The formation of A and B compartments is thought to be driven by a combination of factors including the distribution of active and repressive chromatin marks (3). At mid-range distances, topological domain structure is formed by high density of contacts inside the same TAD and relatively high insulation between adjacent TADs (4). TADs are suggested to be main functional regulatory domains by modulating contacts between enhancers and promoters. Elevated frequency of contacts within the TAD mediates a physical interaction between enhancer-promoter pairs, while high insulation at TAD borders can restrict such interactions for pairs located in neighboring TADs (5). At the finest scale, chromatin is organized into loops that bring promoters close to their regulatory elements such as enhancers. Chromatin loops are thought to be formed through the cohesin-mediated loop extrusion restricted by CTCF and other factors (6–9).

Distinct mechanisms facilitate chromatin feature formation at multiple levels. Besides conventional types described above, the current research (10–17) has revealed an additional level of interactions mediated by Polycomb group (PcG) proteins. PcG proteins are a family of transcriptional repressors that have first been described as capable of gene silencing maintenance during development and cellular differentiation. PcG proteins are organized into two epigenetic complexes, the Polycomb Repressive Complex 1 and 2 (PRC1 and PRC2). PRC2 catalyzes the trimethylation of lysine residue on histone H3 (H3K27me3), which contributes to chromatin compaction and provides a stable and heritable mark that can be passed through cell generations (18, 19). PRC1 is responsible for maintaining the silenced state of genes by recognizing and binding to the H3K27me3 modification produced by PRC2 (14, 20). While the role of PcG proteins has been mostly investigated in the context of the development, their involvement in the maintenance of chromatin organization in mature cells is only starting to be elucidated. Specifically, PcG proteins mediate contacts between H3K27me3-enriched regions located up to hundreds of megabases apart (10, 13–15, 17). The formation of extreme long-range *cis-* and even *trans-*interactions indicates that PcG-mediated contacts represent a distinct hierarchical level of the 3D genome. Whereas the general function of long-range PcG-mediated interactions remains unclear, their dynamic reorganization is found to be important during neural differentiation in mice (13).

Neurons represent a fundamental component of the nervous system. They are characterized by complex morphological and functional properties that require the establishment of specific gene expression programs (21). However, the complexity of the brain tissue and limitation of sample availability constraint the range of conducted chromatin studies focused on main types of brain cells. Few studies compared genome-wide neuronal chromatin organization with non-neuronal cells but they unveiled the presence of neuronal-specific chromatin features at multiple levels of the genome organization (22–24).

Altogether, though some unique features of chromatin organization in neurons have been previously shown, a comprehensive comparison of the chromatin architecture in neurons and non-neuronal cell types is still lacking. Moreover, none of the previous works focus on long-range Polycomb interactions as a distinct feature of the neuronal 3D genome. Here, we use fluorescence-activated nuclear sorting (FANS) to separate neurons and non-neuronal cells obtained from the Wernicke’s area (BA22p) of four human postmortal brain samples, and apply an optimized Hi-C protocol (25) to construct chromatin interaction maps for neuronal and non-neuronal cells. We observe a prominent decrease of chromatin compartmentalization in neurons accompanied by the presence of neuron-specific long-range cis and trans contacts between H3K27me3-marked loci. These long-range interactions could shed a new light on the involvement of PcG proteins in the 3D organization of mature genomes in the brain.

## MATERIALS AND METHODS

### Tissue collection

This study was approved by the Bioethical Commission of the Institute of Gene Biology, Russian Academy of Sciences and was conducted in accordance with the Declaration of Helsinki. Post-mortem human brain samples were obtained via National BioService Russian Biospecimen CRO (St. Petersburg, Russia). Informed consent for using human tissues for research was obtained in writing form from all donors or their next of kin by the tissue provider bank. Sampled brain tissue of all subjects was defined as healthy by medical pathologists. All subjects had sudden death with no prolonged agony state and had no history of neurologic and psychiatric disorders or alcohol and drug abuse. Post-mortem brain frontal blocks 1-1,5 cm thick were stored at -80°C warped in a foil, in zip-lock packages. For sample dissection, The Atlas of the Human Brain (AHB) (26) was used to locate the area of interest, a posterior part of the superior temporal gyrus of the left hemisphere (posterior Brodmann area 22 (BA22p), Wernicke’s area). Frontal blocks with surfaces nearest to AHB level: 60 (MNI: -36.57; ICL: 34.82; MCP: 20.08) were placed into the cryostat Leica CM1950 chamber with -20°C before dissection for 15-20 min on custom-made aluminum support. A tissue piece of approximately 150 mg weight was cut out from a selected area using a metal scalpel. Dissected samples were then collected with tweezers, put into tubes, and placed in dry ice immediately. All materials used during dissection (scalpels, tweezers, tubes) were sterile and chilled using dry ice before use.

### Nuclei isolation

Nuclei isolation from brain tissue and further NeuN antibody immunostaining were performed according to the previously published STAR protocol (25) with optimization for human brain tissue. Briefly, we dissected and fixed 200-300 mg of frozen brain tissue with a 1% formaldehyde-based buffer and homogenized it with a plastic pestle on ice. The brain homogenate was mixed for 10 minutes at room temperature, then the crosslinking reaction was quenched with 2M Glycine and centrifuged at 1100g +4 °C during 5 min. After that, the pellet was washed several times with NF1 buffer (Tris-HCl 10 mM, pH 8.0; EDTA 1 mM; MgCl2 5 mM; Sucrose 100 mM; Triton X100 0.5%), homogenized with Dounce homogenizer 10–30 times on ice and filtered to the new tube. Then sucrose mix (1.2M Sucrose) was layered on homogenized samples to isolate nuclei with ultracentrifugation in a sucrose gradient (1600g, 5 min, +4 °C). The nuclei pellet was resuspended in PBST with 5% BSA and 3% bovine serum, the isolated nuclei were stained with NeuN antibody (1:10000 anti-NeuN-525, FITC) for at least 1 hour, and then resuspended in PBST with 5% BSA. Before FANS (Fluorescence-Activated Sorting of fixed Nuclei) procedure nuclei were filtered with a 35-μm cell strainer into the Falcon™ Round-Bottom Polystyrene Test Tubes.

The sorted NeuN-positive and NeuN-negative nuclei populations were collected and used in amounts of 200,000-400,000 sorted nuclei per sample for Hi-C libraries preparation.

### Hi-C library preparation

Preparation of bulk Hi-C libraries from 8 samples of sorted nuclei from the human brain tissue was performed as previously described (2, 27) with optimal changes to brain tissue (25). Approximately 200,000-400,000 sorted nuclei for the sample were lysed in a solution of lysis buffer containing 150 mM Tris-HCI pH7.5-8, 150 mM NaCI, 0.5% (v/v) NP-40, 1% (v/v), Triton-X100, 1x Protease Inhibitor (CALBIOCHEM, #539137) on ice for 15 min, centrifuged at +4 °C 5000g for 7 min, the pellet was resuspended and washed in 1.12Х restriction enzyme DPN II buffer. After the final wash, the pellet suspension in 1.12Х DPN II buffer with added 0.3% SDS has incubated for 60 min 1400 rpm at +37°C. Further, the reaction was quenched with 1.8% Triton X-100 for 60 min at 1400 rpm at +37°C, and the 500U DPN II restriction enzyme was added, the suspension was left overnight at 1400 rpm at +37°C. After that, the additional amount of 200U DPN II restriction enzyme was added to the digested nuclei and incubated for the next 120 min at 1400 rpm at +37°C. After that, the restriction was unactivated at +65°C for 20 min. The digested DNA samples were washed and resuspended in 1.2XNEB2, and after that digested ends were marked with biotin during the incubation in the mix of 1М biotin-14dATP, 10x NEB2, 10mM dCTP, 10mM dGTP, 10mM dTTP, 5U/ul Klenow (NEB #M0210S) for 90 min 900 rpm at +37 °C. The resulting blunt ends were washed with 1X T4 DNA ligase buffer and chromatin fragments were proximally ligated with 50 U T4 DNA ligase (Fermentas) for 6 h 1400 rpm at +37 °C. Then the 0.5% SDS with 20 μg/ml proteinase K was added to samples for overnight incubation at +65 °C. The following step was DNA purification with phenol–chloroform extraction and precipitation with 2.5 volumes of 96% ethanol, 0.1 volumes of 3M sodium acetate, 20 μg/ml glycogen, and 9.4 μg/ml tRNA for 60 min at −80 °C. After the centrifugation step at 20000g for 20 min, +4°C the pellet was dissolved in 10 mM Tris-HCl pH 8.0 with the addition of RNase A and incubated for 45 min at +37°C. The next step was DNA sample purification on 2.5 volumes of magnetic AMPure XP beads with the following DNA elution in 10 mM Tris-HCl for 15 min at +55°C. Eluted DNA was fragmented by sonication with Power 15 regime, 4 repeats for 30 s with 3 min break. Sonicated DNA samples were then concentrated on Amicon columns 14000g/5 min with double 10 10 mM Tris-HCl pH 8.0 washing. The resulting biotin-marked fragments were isolated using Dynabeads MyOne Streptavidin C1 (Invitrogen #65001) and diluted in TWB buffer (1M Tris-HCl pH 8.0, 50 mM EDTA, 5M NaCl, 1% Tween). All subsequent steps were performed on that bead-bound DNA fraction including DNA ends repair with repair mix (10Х ligase NEB buffer, 10 mM dNTP mix, T4 DNA Pol, T4 PNK, 1 ul Klenow) for 30 min at room temperature, and PolyA tail adding reaction with a mix containing 10Х NEB2, 10 mМ dATP, Klenow (exo-) and incubation for 30 min at +37°C and then for 20 min at +65°C. Then beads were washed with 1X T4 ligase buffer for 3 min at room temperature 900 rpm, and the ligation reaction was performed with a ligation mix of 10X T4 ligase buffer, adapter, 5 U T4 ligase (Fermentas) for 2.5 h at room temperature. Finally, libraries were PCR amplified using Illumina forward and reverse primers and KAPA HiFi HotStart for 16 cycles to select the appropriate cycle for amplified DNA fragments collection (the 6, 9, 12, and 15 cycles were collected and checked for DNA quantity on chosen cycle). Last, up to the chosen number of cycles Hi-C libraries were PCR amplified, and resulting fragments were purified using 1.5 volumes AMPure XP magnetic beads.

### Hi-C data processing

Raw reads of each Hi-C library were trimmed with Trim Galore by setting the parameter *stringency* equal to 3 and then mapped to the human hg38 reference genome with the distiller-nf pipeline (https://github.com/open2c/distiller-nf, v.0.3.3) (28) which utilizes BWA for genome mapping procedure, ensuring valid reads filtering, ‘binning’, and generating list of valid pair contacts which transformed into binned matrices of Hi-C interactions.

Most of the Hi-C analysis was performed on Hi-C matrices merged by cell type. Merging was done at 1 kbp resolution using cooler.merge_coolers() function of a cooler library (29), v.0.8.11. The resulting merged Hi-C maps were sampled to the equal number of contact pairs using cooltools.sample() function of a cooltools library (30), v.0.5.1. To avoid possible bias due to Hi-C ligation artifacts, the values at the main diagonal were removed prior to sampling using cooler and custom python code. Resulting .cool files with Hi-C matrices were ICE normalized (31) using “cooler balance” command with default parameters.

For the reanalysis of the data from Hu et al. (22), non-neuronal sample “HSB181neg” was ignored due to low chromatin structure similarity with other non-neuronal samples from the same study (data not shown).

### Hi-C data quality control

To demonstrate that the chromatin architecture is preserved within the range of post-mortem intervals of the brain samples, we conducted a comparison between mouse neuronal Hi-C data obtained from freshly euthanized (n=2) animals and those from fresh-frozen (n=3) specimens (GSE168524 and GSE172228 datasets, respectively). Additionally, we included the only available glial Hi-C dataset obtained from freshly euthanized (n=2) mice as a reference (GSE168524 dataset). To minimize potential biases, we retained only male replicates for further analysis. The comparison was based on a Stratum-adjusted Correlation Coefficient (SCC), calculated for 20-kb Hi-C maps after downsampling to an equal total number of contacts. The analysis revealed an expected correlation between neuronal samples and their distinct separation from glial ones, indicating a high reproducibility within cells of the same type despite variations in sample preparation strategies and source brain regions (Supplementary Figure S1). This observation supports the assumption that there is no major variation in Hi-C data quality between fresh and fresh-frozen samples.

Furthermore, a previous study examining changes of key epigenetic marks over post-mortem intervals of 0, 24, 48, and 72 hours in large neurons and other brain cells from neonatal pig cortex (32) reported significant differences beginning to manifest only at the 48-hour delay in a few histone modifications. Importantly, our samples exhibited clear separation into NeuN(+) and NeuN(-) groups without notable variation in the distance between cell types across samples (Supplementary Figure S2), despite post-mortem intervals ranging from 5 to 30 hours. The observation suggests that our samples remain preserved to a similar extent within this range of post-mortem intervals.

Taken together, these considerations, along with the analysis of mouse samples from an independent study, collectively indicate the appropriate and comparable preservation of chromatin architecture in both neuronal and non-neuronal cell types.

### Hi-C compartments

Merged Hi-C maps in .cool format (29) at 50 kbp resolution were filtered to remove consecutive genomic regions (bins) that are empty in at least one of two matrices. Hi-C compartments were called using cooltools software (v0.5.1) (30), eigs_cis() function with parameters: n_eigs=5, sort_metric=’spearmanr’. ChIP-seq H3K27ac histone abundance profile was used as a phasing track. Chromosomes Y and M and short chromosomal arms: chr13_p, chr14_p, chr15_p, chr21_p, chr22_p - were ignored. For neurons, the principal component (PC) that correlated best with H3K27ac histone abundance was used to analyze compartments. For non-neurons, we visually compared the first two PCs that correlated best with H3K27ac and for each chromosomal arm chose the PC that best described the Hi-C matrix. Saddle plots were calculated using cooltools.saddle() function with parameters: contact_type=’cis’, n_bins=50, qrange=(0.025, 0.975), min_diag=4. To illustrate that the finer compartment structure was more pronounced in NeuN(-) maps compared to NeuN(+), we used NeuN(-) PC to construct both NeuN(+) and NeuN(-) saddle plots. The code for saddle plots was adapted from cooltools tutorial: https://cooltools.readthedocs.io/en/latest/notebooks/compartments_and_saddles.html. To calculate compartment interaction strength at Figure 1I, the mean intensity of the top 20% of corresponding interactions obtained from the saddle matrix were used.

**Figure 1.**
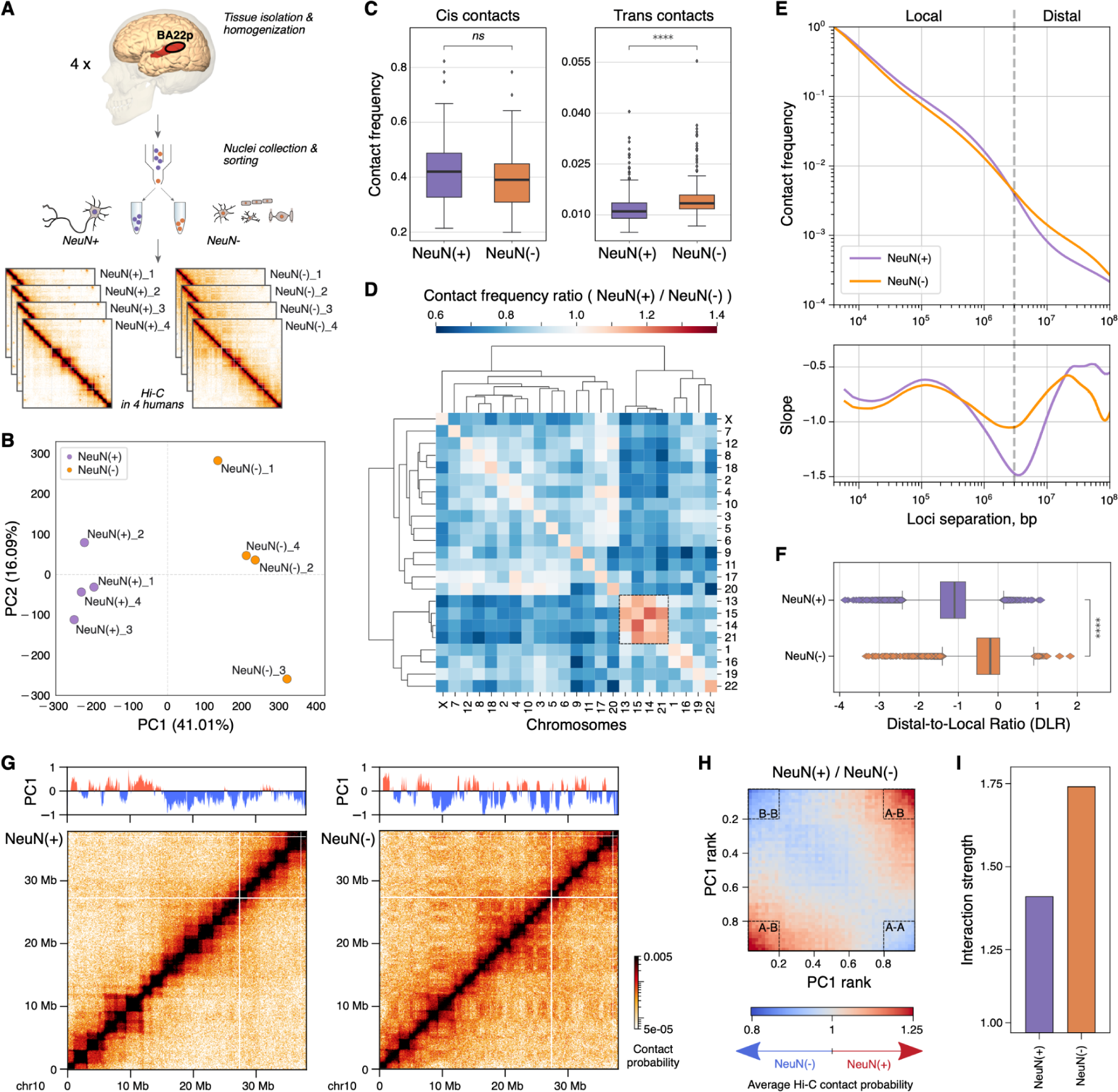
Global chromatin organization in NeuN(+) and NeuN(-) cells. **(A)** Anatomical localization of the analyzed brain region, experimental procedure and the design of this study. Hi-C experiments were performed in NeuN(+) and NeuN(-) cells isolated using FANS from the Wernicke’s speech area of four human individuals. **(B)** Principal component analysis plot based on the Insulation Score (IS) variation among all produced Hi-C maps. Colors represent NeuN(+) and NeuN(-) cells, here and in panels C-F. **(C)** Average interactions within all chromosomes (cis contacts, left panel) and between all pairs of chromosomes (trans contacts, right panel) calculated separately for NeuN(+) and NeuN(-) cells. Asterisks indicate Wilcoxon test p-values: **** - p < 0.00001, ns - p > 0.05. **(D)** Ratio of interactions within and between all chromosomes (NeuN(+)/NeuN(-)). **(E)** Polymer scaling plot showing average interaction frequencies at various genomic distances, as well as the first derivative of this plot demonstrating the slope. **(F)** Distal (>3 Mb) to local (<3 Mb) contact ratio (DLR) calculated for every 100-kb genomic region, demonstrating that chromosomes in neurons are more compact at a local scale and less compact at a large scale. Asterisks represent Wilcoxon test p-value < 0.00001. **(G)** A fragment of the Hi-C map featuring the compartment eigenvector (PC1) for NeuN(+) (left) and NeuN(-) (right) cells. Positive PC1 values represent the A compartment, while negative values correspond to the B compartment. **(H)** NeuN(+) / NeuN(-) ratio of average Hi-C contact probability (observed over expected) of genomic regions arranged by the corresponding PC1 rank (saddle plot). Both saddle plots were created using NeuN(-) PC1. **(I)** Compartment interaction strength calculated as the highest intra-compartment interactions divided by the lowest inter-compartment interactions.

To calculate correlations of Hi-C maps PC1 with chromatin states, the genome was split into 100 kbp bins. Chromatin states annotation for neuronal and non-neuronal cells was retrieved from Dong et al. (33). The annotation includes six distinct chromatin states: ‘Quies’, ‘ReprPC’, ‘TssBiv’, ‘TssFlnk’, ‘TssA’, and ‘EnhA’. For each genomic bin, the proportion of the bin occupied by each chromatin state was calculated. Obtained values of chromatin state proportions were used to calculate pairwise Spearman’s rank correlation coefficients between states and PC1 values. The correlations between PC1 and chromatin states were calculated for neurons and non-neurons separately.

### Insulation score profiling and TAD calling

Insulation score (IS) was calculated genome-wide for 15-kb resolution NeuN(+) and NeuN(-) Hi-C maps with the following parameters: min_frac_valid_pixels=0.82, min_dist_bad_bin=10, window = 120_000. Principal component analysis was based on the obtained IS profile variation among all produced Hi-C maps, as well as similarly re-analyzed publicly available NeuN(+), NeuN(-) Hi-C maps (22) and other tissue and cell line Hi-C maps (34, 35).

The identification of TAD relied on the genome-wide insulation score identification. TAD calling on Hi-C data was implemented with cooltools software (v0.5.2) (30). Module “insulation” available in the software utilizes one of the common algorithms for insulation profile calculation that is based on diamond-window score. Insulation score was defined genome-wide on NeuN(+) and NeuN(-) maps at 15 kbp resolution with the following parameters: min_frac_valid_pixels=0.82, min_dist_bad_bin=10, window=120_000. TAD boundaries were defined based on the border strength with the application of the “Li” threshold from scikit-image (36).

NeuN(+) and NeuN(-) Insulation tables retrieved from cooltools were converted to bedGraph format by extracting columns representing chromosome, start or end of the interval and logarithm of the insulation score. Then, bedGraphToBigWig v.4 binaries available at UCSC Genome Browser web-page were applied to convert insulation score files into bigWig format (37).

### Identification of cell-type-specific TAD borders

Cell-type-specific TADs were defined based on one of the following criteria: 1 - TAD border has no intersection with all other TADs identified in another type of cell, 2 - ratio of border strengths between NeuN(+) and NeuN(-) cells is not within Q1 to Q3. Identification of the cell-specific TAD borders relied on several steps implemented with the bedtools package v.2.30.0 (38). First, each TAD was widened by 1.5 bins (22.5 kbp) at both sides from the original TAD border with “bedtools slop -b 22500”. This step ensures that TAD borders error calculation will not significantly affect the intersection and precision of differential TADs identification. Then, cell-specific .bed files with increased TAD borders were intersected with “bedtools intersect -a glia1.bed -b glia2.bed > all_glia.bed*”* to account for the possible variations between samples. These steps resulted in the TAD borders presented in glial and neuronal samples. Finally, these files were subtracted from each other to identify differential TAD borders with “bedtools intersect -a all_glia.bed -b all_neuron.bed -v > glial_unique.bed”. As the results of the procedure, part of the neuron-specific and glia-specific TAD borders were identified along with common TADs.

Common TADs were inspected further to identify borders whose strength is altered significantly between cell types. First, the ratio of border strength was calculated for each individual border and the following identification of the quartiles of ratio distribution. Borders with ratio absolute value more than third quartile were added to respective cell-specific TAD borders.

### Analysis of TAD covered with LADs

LaminB1 ChIP-seq data from the developing human forebrain (gestation week 20) (39) was used as a proxy for LADs positions. The genomic coordinates of laminB1 peaks were intersected with the positions of TADs in NeuN(+) and NeuN(-). TADs that intersected with laminB1 peaks, covering more than 99% of their size, were categorized as being located within LADs. Conversely, TADs with 0% coverage were classified as positioned outside of LADs.

Size groups of TADs were identified based on the distribution of TAD sizes across all samples. Specifically, TADs were categorized into three distinct groups based on their sizes in relation to the lower and upper one-third quantiles of the distribution.

### Calculation of the TAD density

TAD density was calculated as the average interaction frequency within topologically associating domains (TADs). Specifically, we first normalize the Hi-C contact matrices and then compute the mean interaction frequency for each TAD. Next, we calculate the mean interaction frequency at increasing distances from the diagonal, which represents closer genomic proximity, and normalize the entire matrix by these mean values. This step results in a symmetric, normalized contact matrix for the specified chromosome. Then, we extract the corresponding submatrix from the normalized contact matrix for each TAD. Finally, we calculate the mean interaction frequency for the upper triangle of this submatrix, excluding the diagonal to avoid self-interactions. This process is iteratively repeated for all TADs and chromosomes to provide a quantitative measure of the strength of chromatin interactions within TADs.

### Annotation of loops

Loops positions were identified with the “cooltools.dots” module of cooltools software (v0.5.2)(30). Hi-C maps with the resolution of 5 kbp were utilized. The following parameters were set for loop calling: max_loci_separation = 10_000_000, max_nans_tolerated = 1, lambda_bin_fdr = 0.05, clustering_radius = 20_000, cluster_filtering = None.

### Loops density identification

The loops union was obtained by gathering loop positions found in NeuN(+) or NeuN(-). Then, off-diagonal pileup was determined individually for each loop with “cooltools.pileup” module of cooltools software (v0.5.2). This module allows to extract chromatin density information with the window centered at the pixel with one anchor as a left loop coordinate and another anchor as a right loop coordinate. Then, the ratio between NeuN(+) and NeN-densities of the central pixels were defined for each loop individually. Distribution of ratios of central pixel densities were used to identify loop groups based on quartiles. Thus, four groups representing loops with central pixel density ratio laying lower Q1 (Group 1), between Q1 and Q2 (Group 2), between Q2 and Q3 (Group 3) or higher Q3 (Group 4) were further examined with different approaches.

### LD score regression analysis in loop anchors

To study the link between these loops and human heritable traits, we applied LD score regression to the anchors of loops in Groups 1 and 4 defined above using the stratified LD score regression method, which allows partitioning of heritability from GWAS summary statistics. We obtained GWAS summary statistics files for a total of 234 traits from either the United Kingdom Biobank (UKBB) or from individual GWAS studies. We followed the LD score regression tutorial available at https://github.com/bulik/ldsc/wiki, using default parameters as suggested. The results are presented as LDSC enrichment p-values, which were adjusted for multiple comparisons with the Benjamini-Hochberg method.

### Loops annotation with differentially expressed genes

Loops containing genes that are differentially expressed (DE) between NeuN(+) and NeuN(-) in their anchors became other types of loop groups. Cell-specific loop positions were intersected with the position of DE genes possessing the respective direction of log2FC. Specifically, loops with log2FC > 0.58 were selected for NeuN(+) loop study, and genes with the opposite expression change were utilized for another cell type. The function “pair_to_bed” of pybedtools v.0.9.0 (40) was utilized for retrieval of overlaps between a .bedpe file and .bed. Thus, two groups were identified for NeuN(+) and NeuN(-) based on the positioning of DE genes: loops containing DE genes in one or both anchors and loops with no intersection with DE genes.

### Chromatin features annotation with chromatin states

Chromatin states annotation for neuronal and non-neuronal cells was retrieved from Dong et al. (33). For annotation of TADs, each border was firstly intersected with the chromatin states data. Then, the percent of coverage with each of six states was calculated. The final results are presented as the log2-normalized ratio of the percent coverage of the selected state found in the border to the genome-wide percent coverage of this state. To study the association of the TAD density with enrichment of active or inactive states, we identified two groups of TADs with the increased ratio of ReprPC or EnhA states. Initially, the upper quartile of the distribution of ReprPC or EnhA states coverages within TAD borders of the corresponding cell type was identified. Then, TADs with exceeding coverage were separated into ReprPC or EnhA groups for further density estimation. Significance of change between groups was assessed with a two-sided Mann-Whitney-Wilcoxon test.

Loops annotation was performed in two modes. First, each loop anchor was annotated independently following the strategy described for TADs. Then, loops were characterized based on the chromatin states assigned to the left and right anchor.

### Analysis of neuronal dots

To analyze neuronal dots, we merged our Hi-C maps with publicly available Hi-C (22). Annotation of neuronal dots was done in two steps. First, we performed manual annotation of bright dot regions on a Hi-C map using HiGlass visualization software (v.1.11.7) (41). Second, we filtered our manual selection to keep a subset of significantly enriched Hi-C interactions. For this, we used FitHiC2 software (v.2.0.8) (42, 43) with parameters: “-r 100000”, “--contactType All”. To calculate the *bias* values, we used the inverse of “weight” values obtained after iterative correction procedure and normalized them to the average of 1:

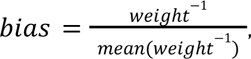

Where *mean* (*weight*^−1^)is calculated over all genomic bins. The resulting FitHiC2 interactions were filtered for q-value < 0.05.

On Figure 4B, FitHiC2 filtering was not used, and all pairs of manually annotated regions were considered. Average interaction was calculated for merged Hi-C maps at 10 kbp resolution using coolpup.py software (v.1.0.0) (44).

Publicly available ChIP-seq data for H3K27me3-enriched regions in ENCODE broadPeak format were used (33).

To find cliques (networks) of NeuN(+) and NeuN(-) interactions, networkx library (v.2.6.3) was used (45). *N* loci form a clique of size *N* if each pair of these loci forms a significant contact according to FitHiC2. Diagrams from Figure 4D were plotted using pyCircos python library (v.0.3.0, https://github.com/ponnhide/pyCircos).

The list of housekeeping genes was obtained from HRT Atlas v1.0 database (46). The list of transcription factors was obtained from Lambert et al. (47).

### snm3C-seq data processing

To study 3D structure in different neural cell types separately, we used publicly available snm3C-seq data (24). To obtain aggregated 3D structure maps, we downloaded “.3C.contact.tsv.gz” processed files (from GSE215353) with contact pairs per cell. Then, using cell metadata table provided by the authors in the manuscript, we selected cells meeting the following criteria:

- <u>for TAD analysis</u>: ‘Region’ = ‘MTG’, ‘MajorType’ = ‘ODC’.
- <u>for neuronal dot analysis</u>: ‘Region’ = ‘MTG’, ‘CellClass’ = ‘Telencephalic excitatory neurons’ or ‘Inhibitory/non-telencephalic neurons’.

For each analysis, we randomly selected an equal number of cells of each cell type and aggregated the contact pairs from the selected cells in contact maps using “cooler cload pairs” command of the cooler library (v.0.8.11) (29). Finally, we applied an iterative correction and eigenvector decomposition procedure to the resulting maps using the “cooler balance” command.

### snm3C-seq data analysis

The annotation of neuronal dots in excitatory (EN) and inhibitory (IN) neurons followed a methodology similar to that used for Hi-C data. We applied the FitHiC2 software (v.2.0.8) (42, 43) with parameters set as “-r 100000”, “--contactType All” for each of the two contact maps and overlapped significant interactions (q-value < 0.05) with the manual dot annotation performed for NeuN(+) Hi-C. Genomic regions that formed at least one neuronal dot in one type of neurons and no dots in the other were classified as forming neuronal dots exclusively in that cell type.

### RNA-seq data processing

Fastq files were processed using nf-core/rnaseq pipeline (48, 49), v.3.8.1, with the following parameters: “--aligner hisat2”, “--genome hg38”, “-profile singularity”. The resulting BAM files were sorted by name using samtools sort (50), v.1.6. Number of read counts per gene was calculated with HTSeq software (51), v.2.0.2, using htseq-count command with parameters: “-t exon”, “--mode intersection-nonempty”, “-s yes” - and GENCODE gene annotation (52), v.41. Protein-coding genes and lncRNAs were further analyzed. Differential expression was calculated using DEseq2 package (53), v.1.36.0. Genes with at least five read counts in at least 5 samples were kept. Two variables: cell type and patient id - were added to DEseq2 design formula. Differentially expressed genes with adjusted p-value < 0.05 and |log2FoldChange| > 0.58 were further used. Gene Ontology enrichment analysis was performed on protein-coding genes using clusterProfiler package (54, 55), v.4.4.4, with enrichGO function and the following parameters: “OrgDb = org.Hs.eg.db”, “pAdjustMethod = “BH””, “pvalueCutoff = 0.05”.

### ChIP-seq data processing

Fastq files were processed using the nf-core/chipseq pipeline (v.2.0.0) (49, 56), with the following parameters: “--macs_gsize 2700000000”, “--genome hg38”, “-profile singularity”.

### Protein extracts preparation and immunoblotting

FANS-sorted nuclei were incubated in RIPA buffer (150 mM NaCl, 1% Triton X-100, 0,5% sodium deoxycholate, 0.1% SDS, 50 mM Tris-HC (pH=8.0), 0,1mM PMSF, and protease inhibitors) for 30 min on ice. The protein extracts then were sonicated at power 15 regime for 10 s with a VirSonic 100 ultrasonic cell disruptor (Virtis). The protein concentration was measured with the Bradford Assay (Sigma-Aldrich). Aliquots of the extracts were separated by sodium dodecyl sulphate-polyacrylamide gel electrophoresis and transferred onto polyvinylidene difluoride membranes (Amersham/GE Healthcare). The membranes were incubated with primary antibodies in a blocking solution (2% BSA in PBS containing 0.1% Tween 20 (PBS-T)) overnight at 4℃. After washing with PBS-T (10 min, three times), the membranes were incubated with secondary anti-rabbit antibodies (horseradish peroxidase-conjugated) for 1 hour at room temperature. The membranes were washed in PBS-T again (10min, five times).

The binding was visualized using a Pierce ECL plus western blotting substrate. Band intensities were quantified using ImageJ software (57).

### Statistical methods

Statistical analyses were carried out using the Python v3.7.4 and R v4.0.2 software packages. Group comparisons of continuous variables were done using the two-sided Wilcoxon rank sum test for unpaired samples and the two-sided Wilcoxon signed rank test for paired samples. Correlation coefficients were determined using Pearson correlation. The one-sided Wilcoxon signed rank test was used to determine whether the median of the sample was greater than the theoretical value.

## RESULTS

### Whole-genome maps of chromatin folding in human neuronal and non-neuronal cells

We applied fluorescence-activated nuclei sorting (FANS) to isolate neuronal and non-neuronal cells from the left posterior superior temporal gyrus (Wernicke’s area - BA22p) based on staining with antibodies against neuron-specific marker NeuN (25) (Supplementary Figure S3). 5-kb resolution Hi-C maps were constructed for the NeuN(+) and NeuN(-) nuclei obtained from four human individuals (Figure 1A, Supplementary Table S1). NeuN(+) and NeuN(-) cells demonstrate a clear separation in the principal component analysis (PCA) of Hi-C maps (41% of the explained variance, Figure 1B), suggesting global differences in chromatin architecture between neuronal and non-neuronal cells. Similar analysis combining our data with publicly available Hi-C maps obtained for a different brain region (22) confirms separation of NeuN(+) and NeuN(-) cells (32% of the explained variance, Supplementary Figure S2A) and serves as an additional quality control of our data. Incorporation of publicly available Hi-C data from other human organs into the PCA analysis reveals that NeuN(+) cells possess the most distinctive chromatin organization (Supplementary Figure S2B), while NeuN(-) cells are located closer to other organs and cell lines. We note that the NeuN(-) cell population is highly heterogeneous as it comprises a mix of endothelial cells, microglia, astrocytes, oligodendrocytes, and other glial cell types. Therefore, NeuN(-) Hi-C maps represent an average picture of non-neuronal cells, and do not reflect chromatin organization in particular glial cell types. Thus, we further focus on NeuN(+)-specific chromatin organization features and use NeuN(-) cell population as a baseline for comparisons.

Among global differences in chromatin organization, the analysis of *cis*- and *trans*-contact frequencies reveals a significant decrease in the neuronal *trans*-contacts compared to the non-neuronal ones (Wilcoxon test p-value < 10^-5^, Figure 1C, Supplementary Figure S4A, B), indicating that chromosome territories in NeuN(+) cells are more pronounced. This observation could be explained by the larger nuclear volume in neurons compared to glial cells (58) (Supplementary Figure S3), potentially providing an additional free space in the nucleus for better spatial segregation of chromosomes. Increased nuclear size could also lead to a global decondensation of chromatin in neurons to occupy the free nuclear space, yet we observe an opposite tendency for cis-contacts (Figure 1C, Supplementary Figure S4A, B). Furthermore, recent studies report that the global chromatin structure is robust to dramatic nuclear volume expansion and contraction, with a striking maintenance of loops, TADs, active and inactive compartments, and even chromosome territories (6, 59). Therefore, we can conclude that the nuclear size alone is unlikely to explain the global differences in genome organization between neuronal and non-neuronal cells. At the same time, in NeuN(+) cells we observe a remarkable increase in contact frequency between acrocentric chromosomes 13, 14, 15 and 21 containing nucleolus organizer regions (NORs) crucial for the nucleolus formation (Figure 1D, Supplementary Figure S4C). This observation suggests that the nucleolus is more pronounced in neurons and is in line with microscopy studies reporting larger well-defined nucleolus as a cytological feature of neurons compared to glial cells (60). Of note, the nucleolus is prominently present in both NPCs, which have highly active ribosomal biogenesis (61), and neurons that are still able to grow despite being non-dividing cells (62).

To further explore chromatin architecture at the chromosomal scale, we analyzed intra-chromosomal contact frequencies at different genomic distances and calculated the contact scaling as the average interaction frequency over distance separating two genomic loci, which can reveal general properties of the chromosome fiber (63). Contact scaling plots show an increase in short-range and a decrease in long-range chromatin interactions in NeuN(+) cells compared to NeuN(-) (Figure 1E), indicating that chromosomes in neurons are more compact at a local scale and less compact at a large scale. To estimate the significance of these differences, we employed a contact-scaling metric calculated separately at every 100 kb region across the genome, the distal (>3 Mbp) to local (<3 Mbp) contact ratio (DLR) (64). A pronounced decrease in DLR (ΔDLR = -0.9) confirms the significance of chromatin interactions shift towards shorter ranges in neurons (Wilcoxon test p-value < 10^-5^, Figure 1F, Supplementary Figure S5). Importantly, these results are consistent with previous studies conducted on a different cortical region (23) and employing a different chromatin capture methodology, namely snm3C-seq (24).

### Weak compartmentalization is a characteristic feature of chromatin in NeuN(+) cells

Remarkably different contact frequency at large genomic distances between NeuN(+) and NeuN(-) cell populations potentially implies different chromatin compartmentalization in these cells. Indeed, both visual comparison and eigendecomposition of Hi-C maps suggests different positioning and weaker chromatin compartmentalization in neurons compared to non-neuronal cells (Figure 1G, Supplementary Figures S6 and S7). Specifically, we observe decreased intra-compartment (A-A and B-B) interactions and increased inter-compartment interactions (A-B) in neurons compared to NeuN(-) cells (Figure 1H, I, Supplementary Figure S8), which is concordant with previous observations (23, 24). Consistently, correlations between NeuN(-) PC1 and NeuN(-) chromatin states are higher than between NeuN(+) PC1 and NeuN(+) states (Supplementary Figure S9). Of note, weaker compartmentalization is observed not only in the temporal cortex in our study but also in publicly available prefrontal cortex Hi-C data (22) (Supplementary Figure S10).

### TAD borders in NeuN(+) cells are enriched with neuron-specific genes

In addition to differences observed at the level of chromatin compartments, visual inspection of Hi-C maps reveals a number of locus-specific differences in TAD structure between neuronal and non-neuronal cells, specifically in a vicinity of genes involved in the neuron-specific processes such as the maintenance of synapse and transport systems, alternative splicing regulation and others (Figure 2A). To analyze TAD profiles in NeuN(+) and NeuN(-) cells systematically, we performed TAD calling using the insulation score (IS) algorithm and identified 2194 NeuN(+)-specific and 2218 NeuN(-)-specific TADs borders, along with 2162 common TADs borders (Figure 2B). We note that, though many TAD borders will be further referred to as cell-specific, they might in fact insulate chromatin in all cell types but with different strength. While there are instances of TAD borders that appear to be unique to NeuN(-) cells (Supplementary Figure S11), we have opted against integrating them into a systematic analysis because of the inherent heterogeneity within the NeuN(-) population. On average, TADs are more pronounced in neurons as compared to non-neuronal cells (Figure 2C, D). Increased contact density at TAD level is in agreement with the enrichment of short-range chromatin interactions (<3 Mbp) in neuronal cells (Figure 1D). However, it should be noted that the weaker TADs observed in non-neuronal cells may be attributed to the heterogeneity of the NeuN(-) population, which could lead to less distinct chromatin features on average.

**Figure 2.**
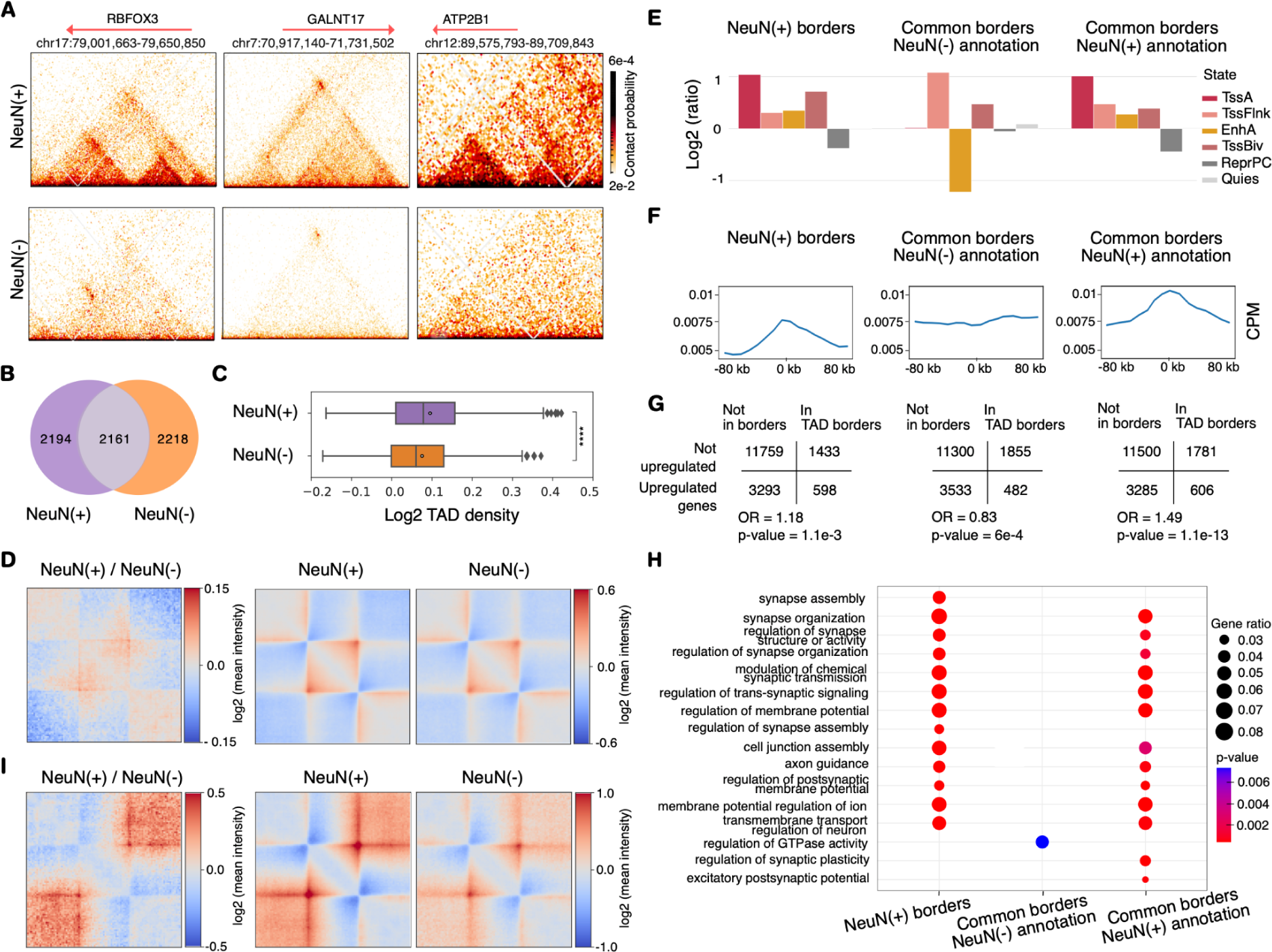
Cell-type-specific differences of TADs in NeuN(+) and NeuN(-) cells. **(A)** Examples of differential TAD profiles at neuronal DE genes. *RBFOX3, GALNT17, ATP2B1* genes location are shown. *RBFOX3* produces the neuronal nuclei (NeuN) antigen that is used as a neuronal marker in the study. NeuN is a RNA binding protein involved in the regulation of alternative splicing of pre-mRNA. *GALNT17* encodes a glycosyltransferase enzyme involved in the post-translational modification of neuronal proteins. This gene has been shown to be essential for the proper development and function of the nervous system. *ATP2B1* is a gene that is involved in neuronal signal transmission encoding a calcium pump that is important for the regulation of calcium levels in neurons. **(B)** Venn diagram of the intersection of neuronal and non-neuronal TAD borders identified based on the insulation profiles. **(C)** Box plot of TAD densities for neurons and non-neuronal cells. Asterisks indicate Wilcoxon test p-values: **** - p < 0.0001. **(D)** Average TAD. **(E)** Grouped stacked bar plots of the chromatin states coverage within TADs borders. The ratio of each state is normalized to the total coverage within the genome in the corresponding cell type. Chromatin states annotation includes six distinct chromatin states, specifically EnhA (active enhancer), TssA (active promoters), TssBiv (bivalent promoters), TssFlnk (promoter flanking region), ReprPC (Polycomb repression region), and Quies (other regions). **(F)** Expression profiles around the cell-type-specific and common borders. **(G)** Confusion tables with Fisher test p-values for the enrichment of genes upregulated in NeuN(-) and NeuN(+) cells within corresponding TAD borders. DE genes were determined as genes upregulated in the selected cell type with FC > 1.5. **(H)** GO terms enrichment (shown with dots sizes) among upregulated genes placed in neuronal-specific or common TAD borders. **(I)** Average large TADs located within lamina-associated domains (LADs).

We therefore proceed with the analysis of epigenetic and structural properties of common and cell-type-specific TAD borders in order to find a connection between TAD border positions and cell-type-specific transcription. First, we examine chromatin states predicted by ChromHMM for NeuN(+) and NeuN(-) cells (33) and find that chromatin states are distributed not equally relative to TAD borders in neurons (Figure 2E). In NeuN(+) cells, both common and neuron-specific TAD borders are significantly enriched with active chromatin states defined by the presence of H3K4me3 and/or H3K27ac histone modifications (TssA, EnhA, TssFlnk), and depleted with repressed chromatin state defined by H3K27me3 (ReprPC).

To verify that the observed differences are not affected by potential inaccuracies in ChromHMM annotations for NeuN(-) cells due to their inherent heterogeneity, we conducted a parallel analysis using chromatin 3D contact maps of oligodendrocytes along with the corresponding H3K27ac/H3K27me3 ChIP-seq tracks (65). For this analysis, we constructed bulk 3D contact maps for oligodendrocytes in the human temporal cortex using publicly available single-nucleus data on chromatin organization (snm3C-seq, see Methods) (24). Consistent with the findings for NeuN(-) cells, we observed a reduction in H3K27ac and a concurrent increase in H3K27me3 coverage around TAD borders for oligodendrocytes (Supplementary Figure S12).

In line with these findings, we observe the increase in mean expression profiles around common and cell-type-specific borders in neurons in contrast to non-neuronal common borders (Figure 2F). Next, we examine the distribution of genes differentially expressed (DE) in NeuN(+) and NeuN(-) cells, according to previously published data (66). In line with active chromatin state profiles, common as well as cell-type-specific TAD borders in neuronal cells are significantly enriched in DE genes upregulated in neurons (Figure 2G) and strongly associated with neuron-specific GO terms (Figure 2H). Remarkably, lists of GO categories are nearly identical for common and NeuN(+)-specific borders. Collectively, these findings suggest the association between TAD borders and transcription in neuronal cells.

We further expand this analysis by categorizing TAD borders into four types, based on the presence of CTCF binding sites and compartment borders. Our findings indicate that in NeuN(+) cells, TAD border characteristics predominantly align with border types associated with CTCF binding sites, rather than compartment borders (Supplementary Figure S13A-D). Conversely, in NeuN(-) cells, compartmentalization appears to have a stronger impact on TAD borders compared to the presence of CTCF binding sites (Supplementary Figure S13E-H).

Furthermore, we segregated TADs into two distinct groups based on their coverage with lamina-associated domains (LADs), that are chromatin regions attached to the nuclear envelope (67). LADs positions were approximated based on the lamin B1 occupancy sites in the developing human forebrain (gestation week 20) (39). It is important to note that some differences in lamin B1 occupancy between the developing and adult human brain are expected, which could potentially interfere with the results. Subsequently, two groups of TADs located outside or within LADs were further classified into three size categories. Remarkably, large neuronal TADs nestled within LADs display distinct characteristics when compared to other TAD groups in NeuN(+) and the corresponding group in NeuN(-) (Figure 2I, Supplementary Figure S14). Specifically, we observe increased interactions at their borders, pronounced extrusion tracks, and increased interactions with surrounding TADs. While these patterns could imply a change in compartmentalization signs, our findings demonstrate that, conversely, large TADs within LADs are consistently surrounded by TADs of the same compartment in NeuN(+) cells (Supplementary Figure S15). Notably, borders of large TADs residing in LADs are enriched with GO terms associated with neuronal development (Supplementary Figure S16). This finding suggests that the positioning of genes related to neuronal development within TADs covered by LADs may be crucial for repressing their expression in mature cells. However, the presence of these genes at the borders could also indicate their conditional activation, potentially occurring during specific developmental stages.

### Loops in NeuN(+) cells facilitate the formation of gene-enhancer hubs

Prominent differences in TAD profile and gene expression patterns between NeuN(+) and NeuN(-) cells imply differential looping between regulatory elements of the genome. Visual inspection of Hi-C maps reveals a number of differential loops, in particular between neuronal enhancers and genes with neuron-specific expression, for example, *ENC1* and *GRIN3A* (Figure 3A). Genome-wide annotation of loops in 5-kb resolution Hi-C maps identifies 1078 shared loops and 1545 NeuN(+)-specific loops (Figure 3B). We observe that the average loop intensity across cell-specific and common loops is the same for NeuN(+) and NeuN(-) cells (Supplementary Figure S17A, B). Notably, neuronal loops are significantly longer than loops in non-neuronal cells (Figure 3C) and, in particular, this trend extends to the subtype of loops associated with CTCF (Supplementary Figure S18).

**Figure 3.**
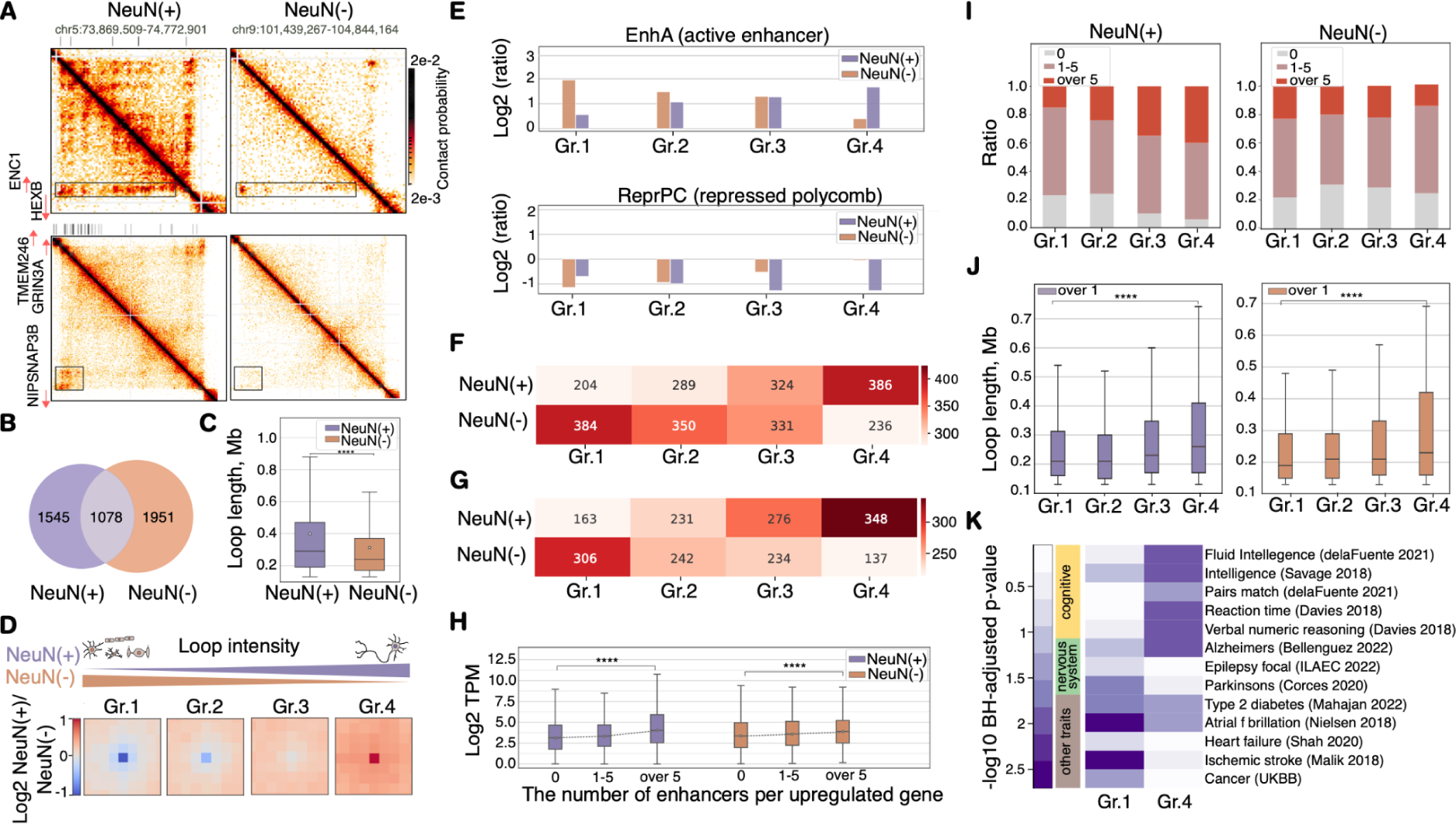
Loops in NeuN(+) and NeuN(-) cells. **(A)** Fragments of 5-kb resolution Hi-C maps for regions of neuronal-specific gene interactions with enhancers. Enhancer tracks are shown at the top of Hi-C maps. **(B)** Venn diagram of the intersection of neuronal and non-neuronal loop positions. **(C)** Box plot of loop length distributions. Asterisks indicate Wilcoxon test p-values: **** - p < 0.00001. **(D)** Average loops within four groups defined based on the increase of NeuN(+)/NeuN(-) intensity ratio. **(E)** EnhA and ReprPC chromatin states in anchors of four loop groups calculated relative to the average across the genome as log2(ratio). **(F)** The number of upregulated genes placed within four loop groups. DE genes were determined as genes upregulated in the selected cell type with FC > 1.5. **(G)** The number of enhancer – promoter loops within four loop groups. **(H)** Box plot of gene expression grouped based on the number of interactions with enhancers (“0” – no interactions, “1-5” and “over 5” – 1-5 or more than 5 interactions). Asterisks indicate Wilcoxon test p-values: **** - p < 0.00001. Groups “0”, “1-5” and “over 5” consist of 3038, 461, 398 and 3227, 496, 295 genes in NeuN(+) and NeuN(-), respectively. **(I)** Ratio of genes separated by three types of interactions with enhancers within four loop groups. **(J)** Box plots of loop lengths mediating interactions between a gene and at least one enhancer. Asterisks indicate Wilcoxon test p-values: **** - p < 0.00001. Gr.1-2, 4 consist of 1227 loops, and Gr.3 consists of 1226 loops. **(K)** Linkage disequilibrium (LD) score regression p-values calculated based on GWAS studies (92–101) for NeuN(-) and NeuN(+) loops. UKBB - UK Biobank (102).

We then grouped shared loops based on their intensity ratio between neuronal and non-neuronal cells and determined groups 1 and 4 as strongest loops in NeuN(-) and NeuN(+) cells (hereinafter these loops are referred to as NeuN(-) and NeuN(+) loops, respectively, Figure 3D). We first annotated chromatin states at loop anchors and observed that NeuN(+) loops are markedly enriched with NeuN(+) active enhancers and depleted with Polycomb-repressed chromatin state defined in NeuN(+) (Figure 3E). Similarly, NeuN(-) loops are enriched with NeuN(-) active enhancers and depleted with NeuN(-) Polycomb-repressed chromatin state. Furthermore, both NeuN(-) and NeuN(+) groups are enriched with DE genes upregulated in NeuN(-) and NeuN(+) cells, respectively (Figure 3F, Supplementary Figure S17C), potentially suggesting the regulatory role of these contacts.

In addition, we show that both NeuN(-) and NeuN(+) loops are enriched with enhancer–promoter interactions (Figure 3G). Moreover, genes interacting with multiple enhancers are on average expressed at a higher level as compared to other genes (Figure 3H), and NeuN(+) loops are enriched with genes interacting with multiple enhancers (Figure 3I). The presence of gene - multiple enhancers interactions especially pronounced in differential neuronal loops could be connected with a wider range of loop lengths (Figure 3J), presumably contributing to establishing interaction on a larger spectrum of distances between genes and enhancers. We thus conclude that the NeuN(+)-specific transcription program might be characterized by the formation of “multivalent” active transcription hubs (68).

Furthermore, we applied Linkage Disequilibrium (LD) score regression to the NeuN(-) and NeuN(+) loops to understand how these loops are related to the human heritable traits (Figure 3K, Supplementary Figure S19). Intriguingly, we find a pronounced and significant enrichment of NeuN(+) loop anchors with cognitive traits exclusively. By contrast, NeuN(-) loop boundaries are enriched with non-brain-related traits, such as autoimmune, blood and cardiovascular disorders, in line with previous studies reporting general enrichment of disease-related genes at loop boundaries (69, 70). Collectively, these findings suggest that NeuN(+)-specific loops might regulate specialized transcription programs associated with human cognitive functions.

### NeuN(+) chromatin is shaped around networks of long-range contacts between H3K27me3 loci

A remarkable feature of the NeuN(+) Hi-C map is the presence of bright dots located far away from the main diagonal and thus representing strong long-range contacts (we further refer to these features as “neuronal dots”, Figure 4A, Supplementary Figure S20). We manually annotated 214 genomic loci that correspond to neuronal dots (Supplementary Table S2) and then mapped significantly high Hi-C interactions of these loci using FitHiC2 software (42) to get neuronal dot annotation. We observe that neuronal dots are longer than regular chromatin loops described in the previous section (Supplementary Figure S21). Furthermore, Hi-C contact frequency of neuronal dots, on average, is much higher compared to contact frequency of the same pairs of regions in NeuN(-) (Figure 4B). Hi-C contact frequency of neuronal dots decays with genomic distance remarkably slower than the genome-average (Figure 4C) pointing at the presence of a molecular mechanism ensuring juxtaposition of dot anchors through large genomic distances. Neuronal dots form networks that include up to 11 loci (Figure 4D). These networks can involve inter-chromosomal interactions, which are present almost exclusively in neurons (Figure 4E). Separate analyses of Hi-C data from the temporal cortex (this study) and prefrontal cortex (22) demonstrate that neuronal dots are present in both regions of the brain cortex (Supplementary Figure S22).

**Figure 4.**
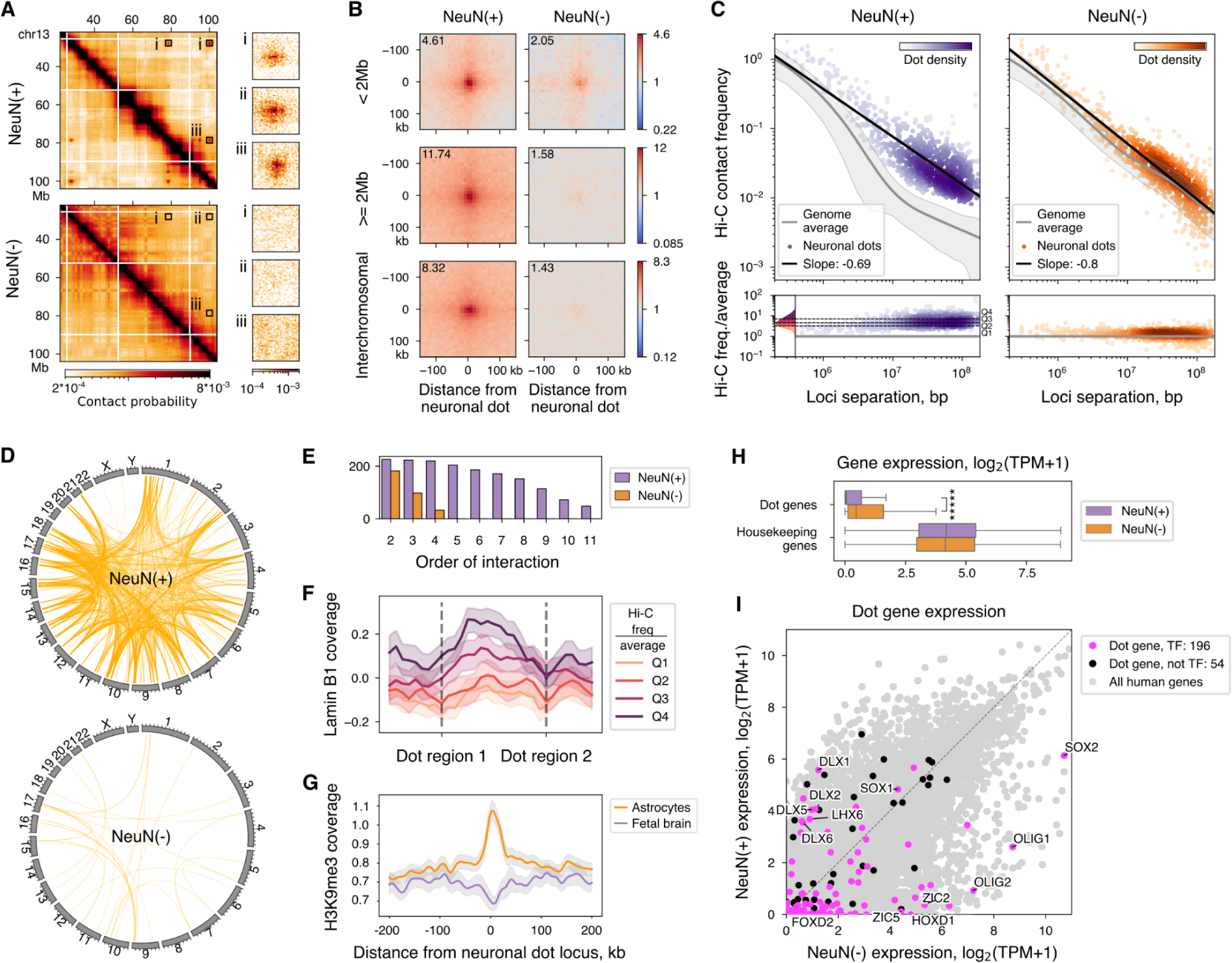
Features of neuronal dots. **(A)** Fragment of the Hi-C map with neuronal dots present in NeuN(+) (top), but absent in NeuN- (bottom). **(B)** The average Hi-C signal (observed over expected) of neuronal dots (left) and corresponding pairs of neuronal dot loci in NeuN(-) (right). The value in the corner corresponds to the central pixel. **(C)** Left: contact scaling of neuronal dots (purple), approximated by a linear regression (black line). Right: contact scaling of pairs of neuronal dot loci in NeuN(-) (orange), approximated by a linear regression (black line). The gray line represents the contact scaling of the whole genome, which is the same as in Figure 1E. **(D)** Significant interchromosomal interactions of neuronal dot loci in NeuN(+) and NeuN(-). Significance is defined by FitHiC2, p < 0.05. **(E)** Statistics for the size (order) of cliques (networks) made by neuronal dots (purple bars) or neuronal dot loci in NeuN(-) (orange bars). **(F)** Average Lamin B1 ChIP-seq read coverage profile between pairs of neuronal dot anchors. Horizontal axis is scaled so that separation between each pair of anchors is constant. Anchor locations are marked by gray dotted lines. Pairs of anchors are separated into four quantiles based on corresponding Hi-C interaction intensity (normalized by expected at given loci separation). Areas around the coloured lines - standard deviations. **(G)** Average H3K9me3 ChIP-seq read coverage profile at neuronal dot anchors. Area around lines - 95% confidence interval. **(H)** Expression of protein-coding genes located within anchors of neuronal dots compared with expression of housekeeping genes. ***** - P-value < 10^-11^, two-sided Wilcoxon test, N=252. **(I)** Expression of protein-coding genes located within anchors of neuronal dots. Genes are divided into TFs (magenta) and non-TFs (black). Grey dots - expression of all human protein-coding genes.

Comparison with available epigenetic profiles (33) reveals that 92% (197 out of 214) of neuronal dot regions overlap with H3K27me3 ChIP-seq peaks. H3K27me3-marked genomic regions are known to be enriched at lamina-associated domain (LAD) boundaries (67). Therefore, we hypothesize that H3K27me3 positioning near lamina increases the probability of neuronal dot regions to get in contact. Analysis of laminB1 occupancy sites in the developing human forebrain (gestation week 20) as a proxy for LADs (39) reveals that the stronger interacting neuronal dot loci have the greater abundance of LADs within genomic regions encompassing these loci (Figure 4F). By contrast, LADs occupancy is strongly depleted at the precise positions of interacting H3K27me3-marked loci. Specifically, 30 neuronal dots are located within LADs, while 71 are expected according to the random control. By contrast, neuronal dots are significantly enriched at LAD boundaries, which are defined as genomic regions spanning 60 kbp around LADs (34 neuronal dots compared with 8 in the random control, chi-square test p-value = 5.2×10^-8^).

To further study the interplay between heterochromatin and neuronal dots, we analyzed ChIP-seq data for H3K9me3, a histone mark of a constitutive heterochromatin. We compared the fetal brain H3K9me3 data as a proxy for NeuN(+) with astrocyte H3K9me3 data as a proxy for NeuN(-) (71). We observed that neuronal dot regions are enriched with H3K9me3 in astrocytes, but not in the fetal brain (Figure 4G). This might indicate that neuronal H3K27me3-marked loci are less occupied by H3K9me3 in NeuN(+) than in NeuN(-), which could have affected the intensity of dot loci.

Most loci brought together by neuronal dots (171 out of 214) reside near the transcription start sites (TSS) of protein-coding genes (Supplementary Figure S23A). Interestingly, the composition of chromatin states in these TSSs is different between NeuN(+) and NeuN(-): in neurons the largest fraction of TSSs is “bivalent”, i.e. occupied by H3K27me3 together with H3K4me3, whereas in non-neurons most TSSs are occupied by H3K27me3 without H3K4me3 (Supplementary Figure S23B). This difference is specific to neuronal dots, since the genome-wide ratio of “bivalent” and “H3K27me3-only” TSSs is similar in NeuN(+) and NeuN(-) (Supplementary Figure S23C).

Out of all protein-coding genes within neuronal dot loci, 80% code for transcription factors (hereby referred to as “dot TFs”). Most dot TFs are repressed in the adult brain in both NeuN(+) and NeuN(-), though in NeuN(+) the expression is significantly lower (Figure 4H, I). Gene Ontology (GO) analysis shows that dot TFs are enriched with biological functions related to the organism development (Supplementary Figure S24). Indeed, some dot TFs belong to gene families known to be involved in organism development, e.g., *HOX* (72), *SOX* (73) and *DLX* (74). Remarkably, the majority (119 out of 197) of dot TFs are associated with the *Nervous System Development* GO term (GO:0007399, hypergeometric test p-value = 3.2×10^-42^).

To further investigate the role of dot TFs, we analyzed the public data for human gene expression during development (75) and found that most dot TFs are expressed in at least one cell type in the developing organism (Supplementary Figure S25). We conclude that dot TFs are expressed during development and must be repressed in the mature brain.

### Genes from human neuronal dots are coupled with PcG proteins in mouse

Neuronal dot regions overlap with H3K27me3, a histone modification that is deposited by Polycomb repressive complex 2 (PRC2) and can be bound by PRC1 (76). Therefore, we hypothesized that H3K27me3 and Polycomb-group (PcG) proteins are involved in the formation of neuronal dots. In mice, H3K27me3-bound loci establish long-range interactions in embryonic stem cells (ES), neural progenitor cells (NPC) and cortical neurons (CN) (13). We thus explore if mouse orthologs of human dot genes are implicated in H3K27me3-coupled long-range interactions. Indeed, many dot genes overlap with the H3K27me3 mark in mice (219, 169 and 192 out of 245 genes for ES, NPC and CN correspondingly) and form long-range interactions.

Moreover, alongside H3K27me3, the mouse orthologs of dot genes are enriched with the ChIP-seq signal of RING1B (Supplementary Figure S26), a component of the PRC1 complex, which is known to be involved in Polycomb-mediated contacts (14). To further investigate the role of RING1B in the repression of genes within neuronal dots, we analyzed a publicly available dataset on RING1A/B knockout in mouse motor neurons (77). We found that RING1A/B knockout leads to the upregulation of 53% (125) of dot genes and downregulation of only 4% (10) of dot genes (Supplementary Figure S27A). This observation suggests that RING1A or RING1B proteins likely contribute to the repression of dot genes in human neurons.

Since the H3K27me3 mark is deposited by the PRC2 complex, we hypothesized that PRC2 might play an important role in neuronal dot formation. Accordingly, PRC2 knockout in mouse striatal neurons leads to upregulation of TFs, many of which can be positively auto-regulated (78). Furthermore, we find that a large fraction of TFs upregulated in PRC2 knockout overlaps with the neuronal dot TFs (Supplementary Figure S27B), suggesting that PRC2 is indeed involved in the dot formation.

Consequently, there is compelling evidence supporting the involvement of PcG proteins, specifically RING1B, in the formation of neuronal dots.

### Five genomic regions form neuronal dots exclusively in excitatory neurons

To gain a deeper our understanding of the neuronal dot phenomenon, we constructed bulk maps of 3D genome structure in excitatory (EN) and inhibitory (IN) neurons of the human temporal cortex, specifically the middle temporal gyrus, using publicly available single-cell data (24) (see Materials and Methods). Our analysis revealed that the majority of neuronal dots are present in both subtypes of neurons (Figure 5A). However, we identified five genomic regions that exclusively form neuronal dots in excitatory neurons (Figure 5B); notably, no IN-specific dot loci were found. These five genomic regions encompass the genes *DLX5, DLX6, SOX2, GAD2, SLC32A1,* and *ARX*.

**Figure 5.**
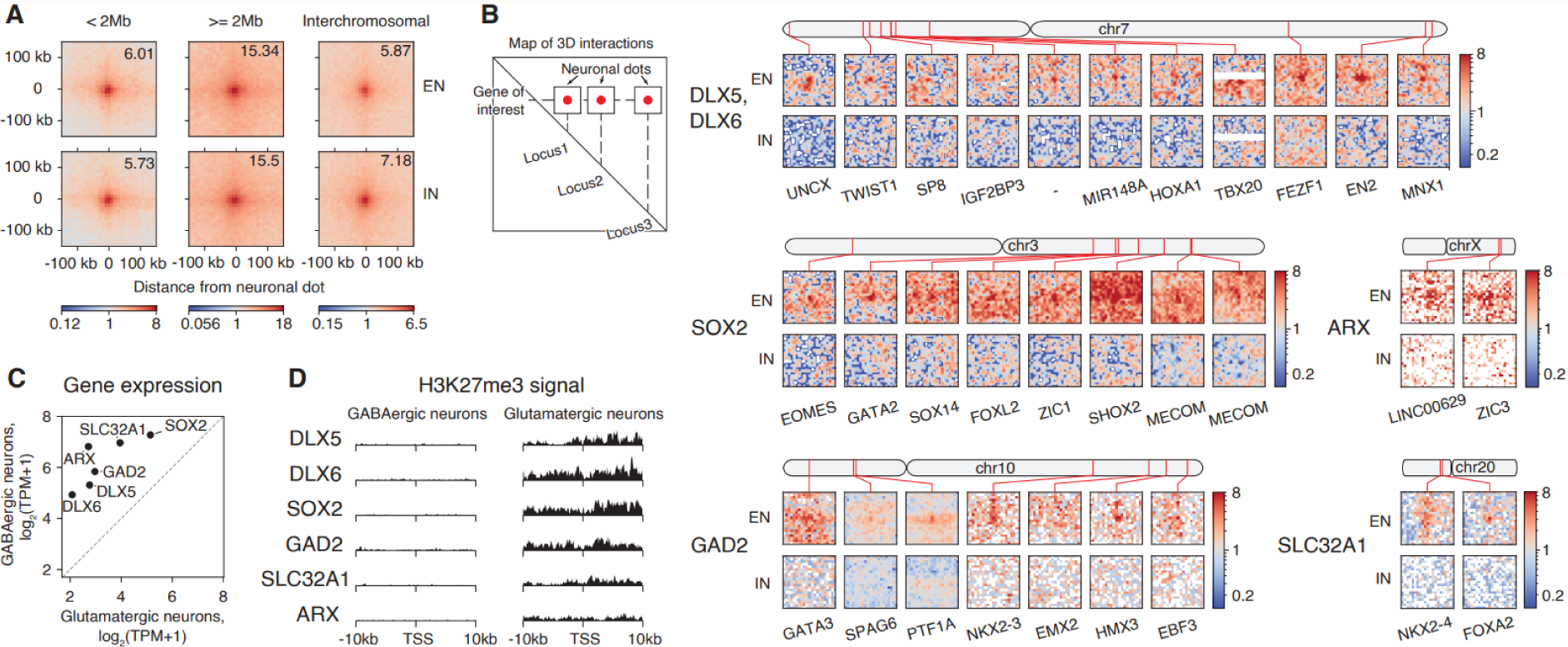
Neuronal dots in excitatory (EN) and inhibitory (IN) neurons. **(A)** Average snm3C-seq signal (observed over expected) of neuronal dots in excitatory (top) and inhibitory (bottom) neuronal dot loci. Value in the corner corresponds to the central pixel. Dot annotation was taken from NeuN(+) Hi-C data. **(B)** Snippets of aggregated snm3C-seq contact matrices showing differential neuronal dots between excitatory and inhibitory neurons. A gene name at the bottom of each plot indicates the gene closest to the dot locus. **(C)** Expression of six genes that form differential neuronal dots. **(D)** H3K27me3 ChIP-seq signal centered at the transcription start sites (TSS) of six genes that form differential neuronal dots. Tracks are oriented so that each gene body is located to the right of the TSS. Vertical axis scale is equal for GABAergic and glutamatergic neurons.

Using publicly available data on gene expression and H3K27me3 mark occupancy in glutamatergic EN and medial ganglionic eminence–derived gamma-aminobutyric acid (GABA)ergic IN (65), we observed notably higher transcription levels of all six genes in IN compared to EN (Figure 5C). Consistently, the H3K27me3 signal around these genes was exclusively detected in EN (Figure 5D). Intriguingly, all six identified genes are known to be involved in the differentiation and/or function of GABAergic neurons. For instance, *GAD2* encodes glutamic acid decarboxylase, crucial for GABA production, while *SLC32A1* encodes the GABA vesicular transporter. *DLX5* and *DLX6* are transcription factors that play roles in the differentiation and function of mature GABAergic neurons (79, 80). *SOX2* is a transcription factor important for neuronal differentiation, although its role in adult neurons is not entirely clear (73, 81). Finally, ARX serves as a marker gene for GABAergic neurons (82).

Thus, subtype-specific expression patterns of genes within neuronal dot loci, coupled with differences in H3K27me3 signal and neuronal dot contact frequency, suggest a potential role of neuronal dots in determining and maintaining neuronal subtype diversity in adult neurons.

## DISCUSSION

Our results suggest that chromatin folding in neurons of the human brain possess several distinct features. First, at the large scale, chromosomes in neurons are more segregated compared to non-neuronal cells as revealed by calculation of a fraction of inter-chromosomal contacts. Intuitively, this observation correlates with the expanded volume of neuronal nuclei (58) as the increase in the nucleus volume provides an additional space for chromatin. Following this logic, individual chromosomes are expected to expand to fill up the free space. Yet, we observe an opposite direction of differences in the total level of intra-chromosomal contacts between NeuN(+) and NeuN(-) cells. Moreover, it has been reported that artificial nuclei expansion (59) does not change global chromatin spatial organization, including chromosome territories. Therefore, factors other than the nuclear volume might contribute to global differences in genome organization between neuronal and non-neuronal cells that we observe.

Neuronal chromosomes are on average more segregated than non-neuronal ones. Notably, chromosomes containing nucleolus organizer regions (NORs) - chromosomes 13, 14, 15, 21, and 22 - display a more intricate interaction pattern. In neurons compared to non-neuronal cells, chromosomes 13, 14, 15, and 21 are closer to each other on average, while chromosome 22 shows no such tendency, indicating differential nucleolar composition in neurons and non-neuronal cells. This observation aligns with previous microscopy studies (83). In particular, Takei et al. applied DNA sequential fluorescence *in situ* hybridization (DNA seqFISH+) technique in mouse brain cells, revealing that the nucleolus has a distinct chromosomal composition in excitatory neurons and astrocytes.

At the local scale, short-range chromatin interactions (< 3 Mbp) are more abundant in neurons. Consistently, TADs are more prominent in neurons, and chromatin loops are, on average, longer. Both neuronal-specific and common TAD borders are enriched with neuron-specific genes and active chromatin marks in neurons. Consequently, we propose that the composition of TAD borders in different brain cell types may align with neuronal transcriptional activity. Potentially, the presence of remnants of neuronal-specific chromatin organization patterns in non-neuronal cells might be attributed to the sequential order of cellular differentiation. Indeed, common precursors first differentiate into neurons, while gliogenesis occurs at a later stage (84). Conforming to the proposed hypothesis, the majority of TAD boundaries exhibit a conservative positioning throughout the developmental transition from ESCs to neural cell types, with only a small fraction of TAD boundaries emerging or mitigating (85).

Distal intra-chromosomal contacts (> 3 Mbp) decay fast in neurons. The slope of contact decay with distance reaches -1.5 in comparison with -1 in non-neurons. This difference cannot be explained by the increased nucleus size in neurons as Sanders et al. demonstrate, on the contrary, slower contact decay with distance upon nuclear volume swelling (59). But they show that increased nucleus size leads to compartment strength weakening, similar to what we and others (23) observe in neuronal Hi-C. However, nuclear swelling does not affect the genomic positioning of compartments (59), while compartments in neurons are largely different from non-neurons.

Schwarzer et al. demonstrated that, in mice, a deficiency of the cohesin-loading factor Nipbl leads to a reduction in TAD strength while enhancing compartmentalization (86). Considering our observations of a similar pattern when comparing non-neuronal cells to neurons, and given that TAD formation is facilitated by the loop extrusion machinery (84, 85), we hypothesize that the differences observed in large- and small-scale chromatin structures may be due to more active loop extrusion in neurons. Although some evidence from Hi-C data supports this hypothesis, further research is needed to directly demonstrate the abundance of cohesin-related proteins in brain cells and their direct impact on chromatin organization.

Our observations highlighting differences in chromatin characteristics at both large and small scales between neuronal and non-neuronal cells align with findings from previous research employing similar methodologies (22, 23). However, our study offers several novel insights. Specifically, we uncover distinct properties of large TADs located within LADs, potentially opening a new avenue for further research. Additionally, in our analysis of chromatin loops, we explore the relationship between loops of varying strengths, with a particular focus on loops facilitating interactions between enhancers and up-regulated genes. Moreover, in contrast to previous studies focusing on the prefrontal cortex, our research extends these observations by analyzing the temporal gyrus area. Notably, among our additional findings, the identification of neuronal dots stands out as a unique discovery, markedly enriching our understanding of chromatin architecture across diverse brain cell types.

A remarkable observation of our Hi-C analysis is the presence of bright long-range interactions between neuronal H3K27me3-enriched regions - neuronal dots - that are formed both in *cis* and in *trans*. To our knowledge, this feature of neuronal 3D chromatin structure has not been demonstrated previously. Despite the rapid decay in the whole-genome expected contact frequency in neurons, neuronal dots are able to establish contacts even at large genomic distances up to 100 Mb. These regions form large interaction networks, comprising up to 11 distant loci.

Within the neuronal dots, a remarkable proportion of genes (about 80%) encode transcription factors (TFs), the majority of which are expressed in neural or non-neural tissues during development but are downregulated in mature NeuN(+) and NeuN(-) cells. Since the H3K27me3 mark is associated with Polycomb repression, we suggest that PcG proteins repress TF genes within the neuronal dots. This hypothesis is supported by two studies conducted in mice (77, 78), and demonstrating that knockouts of PRC2 or RING1A/B proteins in mouse neurons lead to the upregulation of many mouse orthologs of dot genes.

RING1B protein, a member of Polycomb repressive complex 1 (PRC1), is of particular interest since it is required for long-range Polycomb interactions (14). Reanalysis of public data from mouse differentiating neurons (13) shows that: 1) many orthologs of dot genes are occupied by RING1B, 2) some of these RING1B-occupied orthologs form long-range interactions in embryonic stem cells, neural progenitor cells and differentiated neurons. This raises a possibility that RING1B binds to neuronal dot loci and is involved in neuronal dot formation.

Another peculiar feature of neuronal dot loci is that 60% of genes located at their anchors have bivalent promoters, i.e. their transcription start sites (TSSs) are simultaneously occupied by H3K27me3 and H3K4me3. In non-neurons only ∼30% of these genes have bivalent promoters. Such “bivalent” genes in neurons were thoroughly investigated by Ferrai et al., who compared “bivalent” and “H3K27me3-only” genes and demonstrated that upon PRC2 enzymatic inhibition “bivalent” genes get upregulated, while “H3K27me3-only” genes do not (87). Thus, it is possible that “bivalent” repressed genes have more potential for activation than “H3K27me3-only” genes. If this hypothesis is true, then an interesting question for further investigation is why some genes in mature neurons, but not in NeuN(-) cells have this potential to become activated.

Is there a link between neuronal dots and “bivalency” of dot genes? “Bivalent” promoters are known to be occupied by poised RNA polymerase II (RNAP) (87, 88), which is dependent on RING1A and RING1B ubiquitination (88). Thus, both long-range interactions and “bivalency” could be a consequence of RING1A and/or RING1B occupancy at promoters of neuronal dot genes.

Are neuronal dots functionally relevant? Do these interactions affect gene expression at corresponding loci? Kraft et al. demonstrated that long-range PcG-mediated interactions facilitate H3K27me3 spreading, such that the deletion of one interacting locus leads to gene activation in the other (17). While we observe significant downregulation of gene expression at neuronal dot loci compared to NeuN(-), in general, genes within these regions are repressed in both groups. This makes sense, since considered genes are enriched with developmental functions and probably should be inhibited in all types of mature cells.

What biological mechanisms can facilitate such strong PcG-mediated interactions in neurons? One possible explanation is that neurons reside in a post-mitotic state, unlike non-neurons. In general, chromatin organization is largely altered during the cell cycle (89). Ma and Buttitta demonstrated that, in *Drosophila*, H3K27me3 histone mark clustering increases for cells with slower proliferation and reaches maximum for post-mitotic cells (90). They further show that delaying cell cycle exit disrupts H3K27me3 clustering. Thus, a neuronal post-mitotic state might facilitate interactions between H3K27me3-enriched genomic regions.

Interestingly, we observe that the higher abundance of LADs between, but not within, neuronal dot loci correlates with stronger Hi-C interactions. It is possible that chromatin anchoring to nuclear lamina facilitates interactions of the neighboring chromatin regions. An interesting open question is the interplay between PcG-mediated interactions and LADs/compartmentalization. Siegenfeld et al. developed a LIMe-Hi-C experiment that allows to simultaneously capture genome 3D structure and lamina association, and demonstrated that inhibition of EZH2 protein, a catalytic subunit of PRC2, leads to increased compactization of heterochromatin attached to the nuclear lamina (91). They suggested that PcG-mediated interactions and LAD interactions are antagonistic processes. This hypothesis agrees with our NeuN(+) Hi-C data where we observe neuronal dots and reduced B compartment strength compared to NeuN(-).

In conclusion, neuronal genome organization is largely different from non-neuronal one. Many biological processes, including those involved in facilitating interactions over both long and short distances, likely work differently in neurons and probably regulate correct gene expression in this cell type.

## DATA AVAILABILITY

Hi-C data generated in this study are available in the GEO database under the accession number GSE229816. Other analyzed datasets were derived from sources in the public domain. Processed ChIP-seq H3K27me3, GO-CaRT Lamin B1, chromHMM chromatin states, Hi-C of human NeuN(+) and NeuN(-) cells from BA9 cortex area, and RNA-seq data on glutamatergic and medial ganglionic eminence–derived gamma-aminobutyric acid (GABA)ergic neurons data were obtained from the PsychENCODE Consortium at https://doi.org/10.7303/syn25716684, https://doi.org/10.7303/syn25931622, https://doi.org/10.7303/syn21754060, and https://doi.org/10.7303/syn12034263. Processed ChIP-seq H3K9me3 and CTCF data were obtained from ENCODE Portal at https://www.encodeproject.org/, accession numbers are: ENCFF557YDE, ENCFF985OQZ, ENCSR461VHZ, and ENCSR489QDF. RNA-seq data on human NeuN(+) and NeuN(-) cells and snm3C-seq data were obtained from the GEO database, accession numbers GSE96615 and GSE215353. scRNA-seq dataset on the human fetal gene expression was obtained from descartes.brotmanbaty.org, “matrix of normalized gene expression values”. Publicly available dorsolateral prefrontal cortex (DLPFC), lung, spleen, bladder, hESCs and differentiated mouse neurons Hi-C data were obtained from the GEO database, accession numbers GSE87112, GSE52457, and GSE96107. mESCs and mouse NPC Hi-C data were obtained from the 4D Nucleome Data Portal at https://data.4dnucleome.org/, accession numbers 4DNESDXUWBD9 and 4DNESJ9SIAV5. Custom code that was created for data analyses is available at https://github.com/i-pletenev/NeuN_plus_minus_paper.

## SUPPLEMENTARY DATA

Supplementary Data are available at NAR online.

## AUTHOR CONTRIBUTIONS

Ilya A. Pletenev: Formal analysis, Writing—original draft, Writing - review & editing. Maria Bazarevich: Investigation, Formal analysis, Writing - review & editing. Diana R. Zagirova: Formal analysis, Writing—original draft, Writing - review & editing. Anna D. Kononkova: Formal analysis, Writing - review & editing. Alexander V. Cherkasov: Investigation, Formal analysis, Writing - review & editing. Olga I. Efimova: Investigation, Formal analysis, Writing - review & editing. Eugenia A. Tiukacheva: Investigation, Writing - review & editing. Kirill V. Morozov: Formal analysis, Writing - review & editing. Kirill A. Ulianov: Investigation, Writing - review & editing. Dmitriy Komkov: Investigation, Writing - review & editing. Anna V. Tvorogova: Investigation, Writing - review & editing. Vera E. Golimbet: Resources, Formal analysis, Writing - review & editing. Nikolay V. Kondratyev: Resources, Formal analysis, Writing - review & editing. Sergey V. Razin: Investigation, Writing - review & editing. Philipp Khaitovich: Formal analysis, Writing - review & editing. Sergey V. Ulianov: Investigation, Supervision, Writing—original draft, Writing - review & editing. Ekaterina E. Khrameeva: Conceptualization, Formal analysis, Supervision, Writing—original draft, Writing - review & editing.

## ACKNOWLEDGEMENTS

We are grateful to the Core Centrum of IDB RAS and Skoltech BioImaging and Spectroscopy Core Facility for the excellent technical assistance. We thank the Center for Precision Genome Editing and Genetic Technologies for Biomedicine, IGB RAS for the cell sorting. We thank Prof. Mikhail S. Gelfand (Skolkovo Institute of Science and Technology) and Dr. Aleksandra A. Galitsyna for helpful advice, Anastasia Soldatenkova (Skolkovo Institute of Science and Technology) for the help with RNA-seq data processing, Nikita Vaulin (Skolkovo Institute of Science and Technology) for the help with graphical abstract preparation, Maria Molodova (Skolkovo Institute of Science and Technology) for the help with nuclei sorting, Skoltech Genomics Core Facility and Prof. Maria Logacheva (Skolkovo Institute of Science and Technology) for sequencing and quality control of Hi-C libraries, Arkuda HPC cluster provided by the Skolkovo Institute of Science and Technology for computational resources, and National BioService Russian Biospecimen CRO (St. Petersburg, Russia) for providing brain samples. Lamin B1 ChIP-seq, human Hi-C of BA9 cortex area, H3K27me3 ChIP-seq and chromHMM chromatin states data for this publication were obtained from NIMH Repository & Genomics Resource, a centralized national biorepository for genetic studies of psychiatric disorders.

