## Supplemental Figures and Table 1 for "Extensive long-range polycomb interactions and weak compartmentalization are hallmarks of human neuronal 3D genome"

### Supplementary data

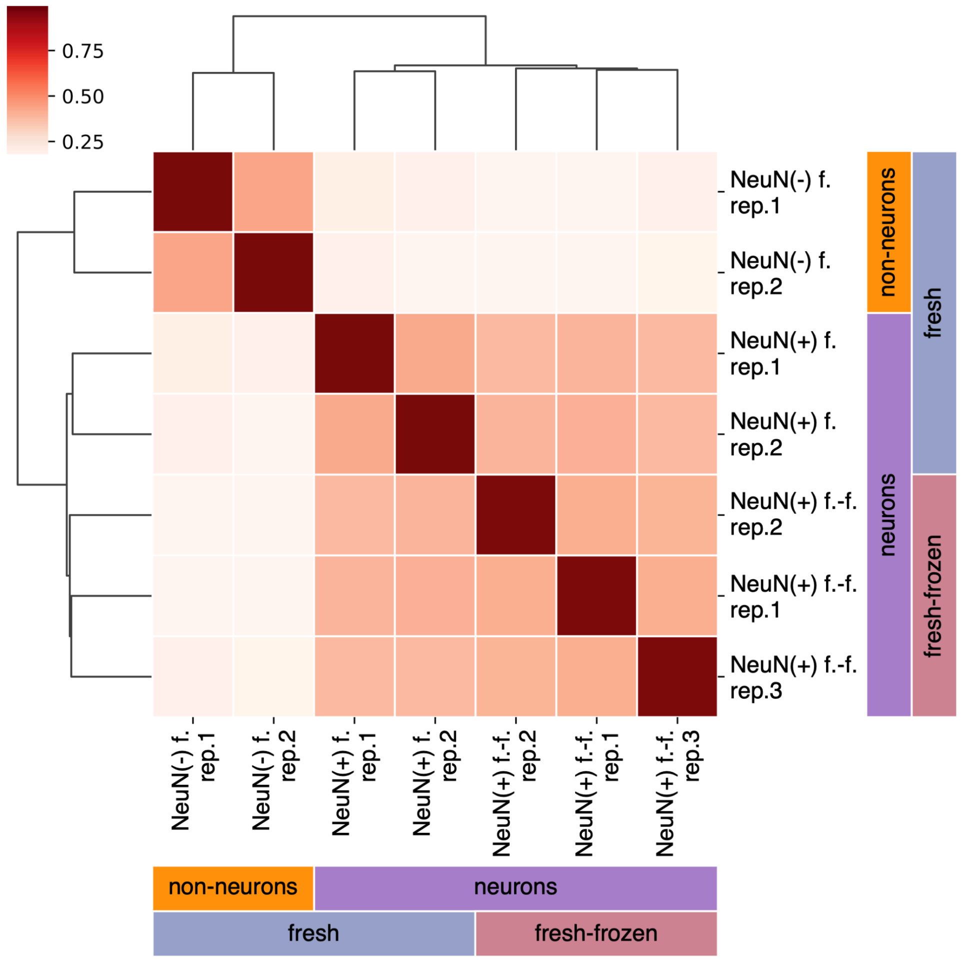


**Supplementary Figure S1**. Clustering of Hi-C maps derived from fresh (f.) and fresh-frozen (f.-f.) mouse samples. The distance between samples was calculated based on the Stratum-adjusted Correlation Coefficient (SCC) value.

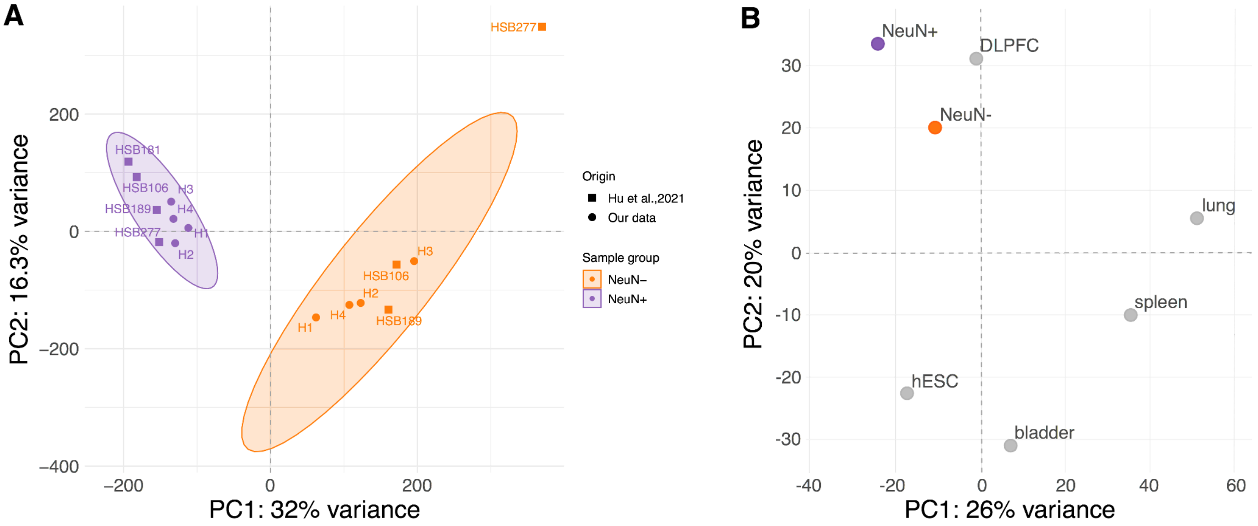


**Supplementary Figure S2.** Principal component analysis plots based on the Insulation Score (IS) variation among our and publicly available Hi-C maps. **(A)** Our data and publicly available NeuN(+) and NeuN(-) Hi-C maps (1). **(B)** Our data and publicly available bulk dorsolateral prefrontal cortex (DLPFC), lung, spleen, bladder Hi-C maps from (2) and hESC - from (3).

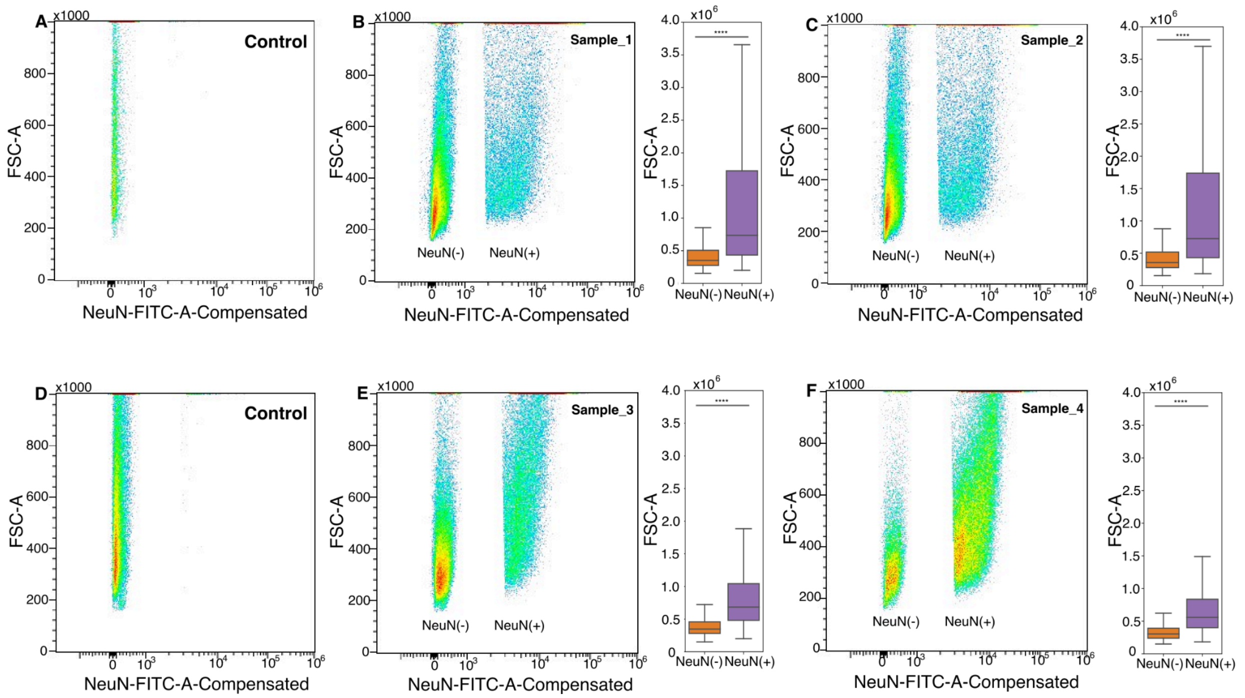


**Supplementary Figure S3.** Fluorescence-activated nuclei sorting (FANS) of NeuN(+) and NeuN(-) samples. **(A-C)** Forward scatter area (FSC-A) vs NeuN signal (FITC-A-Compensated) for Sample_1, Sample_2 and their control sample (autofluorescence). **(D-F)** FSC-A vs FITC-A-Compensated for Sample_3, Sample_4 and their control sample (autofluorescence). Larger values of FSC-A correspond to larger nuclear size. Asterisks above boxplots indicate Mann-Whitney U test p-values: **** - p-value < $10^{-10}$.

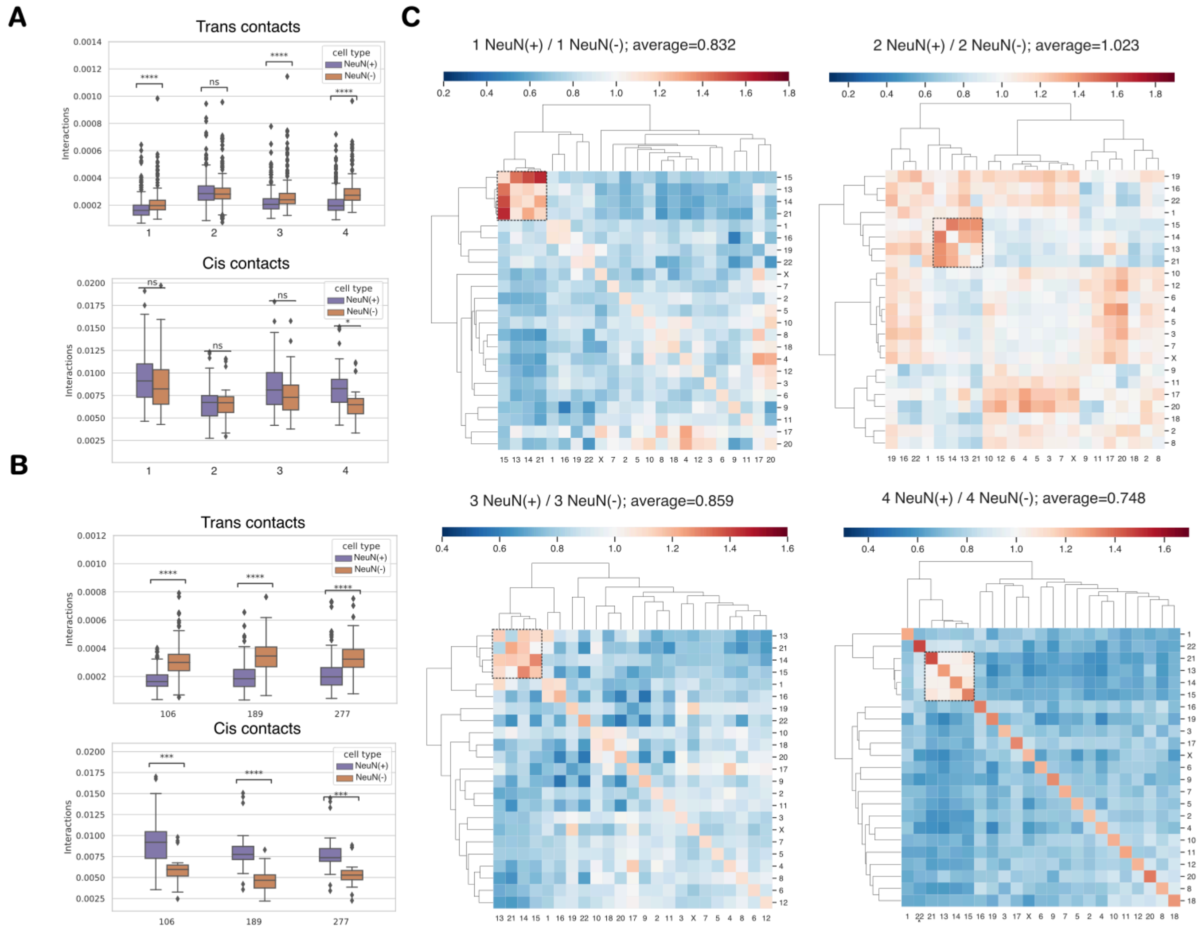


**Supplementary Figure S4**. Interactions within all chromosomes (cis contacts) and between all pairs of chromosomes (trans contacts) in individual brain samples. **(A)** Average trans-contact and cis-contact interactions for NeuN(+) and NeuN(-) cells in four samples from this study. Asterisks indicate Wilcoxon test p-values: **** - p < 0.00001, ns - p > 0.05. **(B)** Average trans-contact and cis-contact interactions for NeuN(+) and NeuN(-) cells in three individual samples from Hu et al. 2021 dataset. **(C)** Ratio of interactions within and between all chromosomes (NeuN(+)/NeuN(-)) for four individual brain samples.

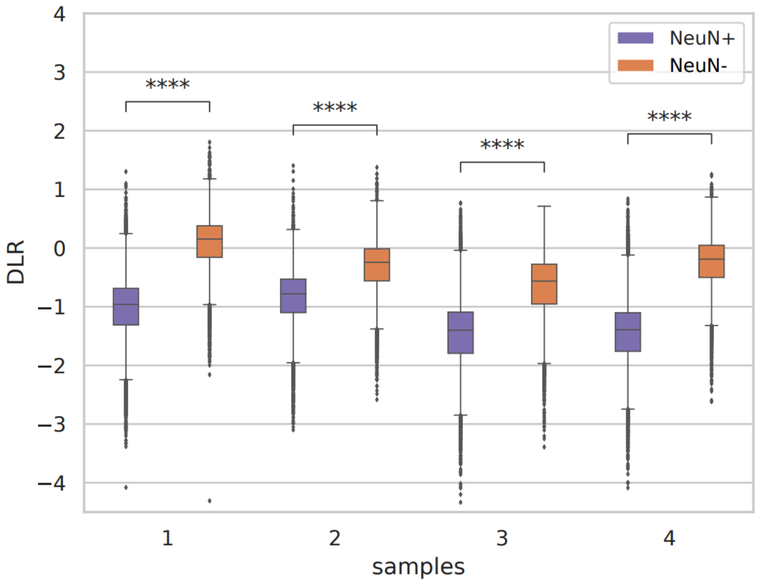


**Supplementary Figure S5.** Distal to local contact ratio (DLR) calculated in four individual brain samples. DLR was calculated for each 250-kb bin as the log2 ratio of the summarized contact frequency with distance separation larger than 3Mb (distal interactions) to the summarized contact frequency with separation less than 3Mb (local interactions).

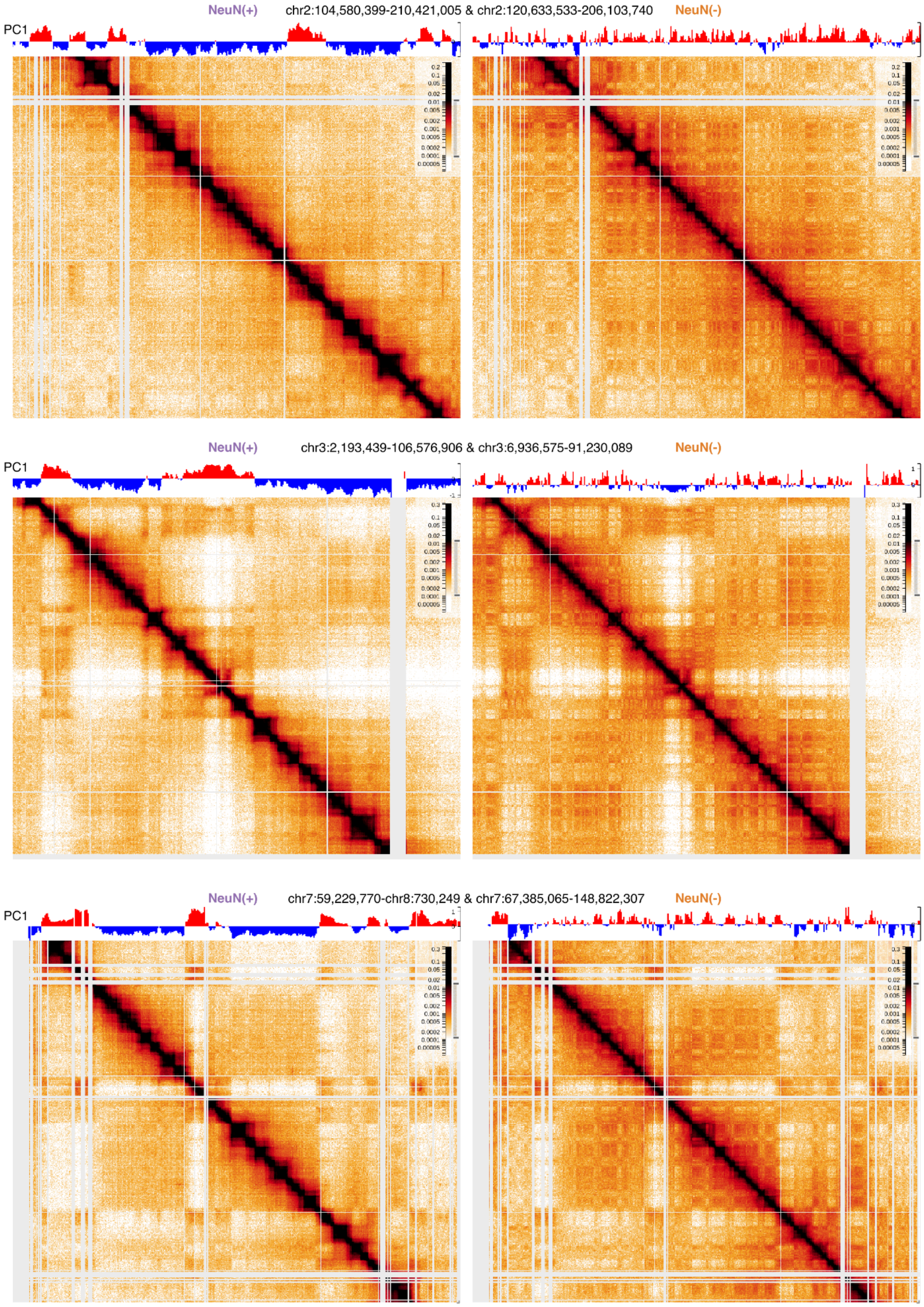


(see legend on the next page)

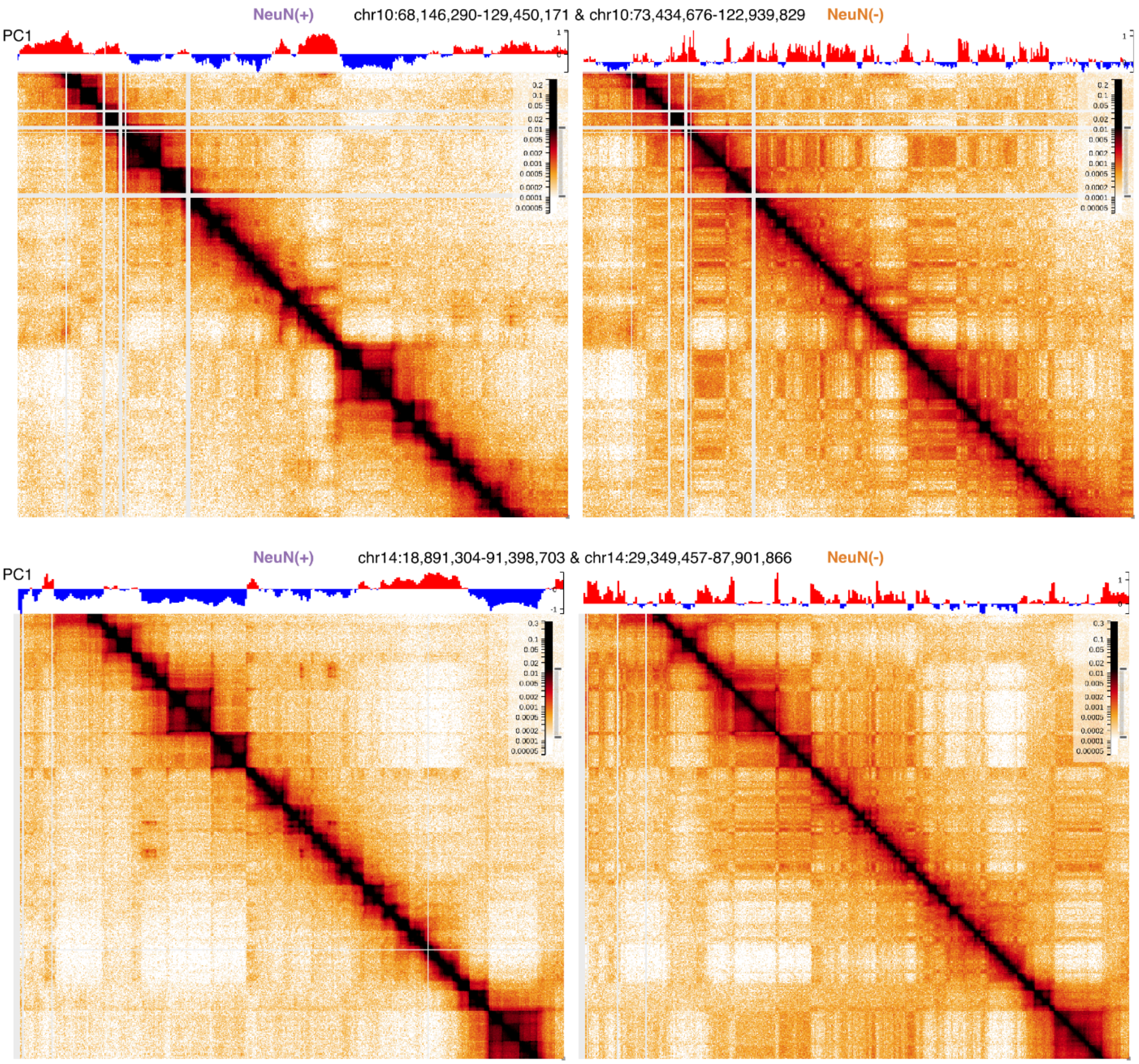


**Supplementary Figure S6.** Examples of different compartmentalization between NeuN(+) (left) and NeuN(-) (right). Above each Hi-C map is the corresponding compartment eigenvector with positive values colored in red (A compartment) and negative values colored in blue (B compartment).

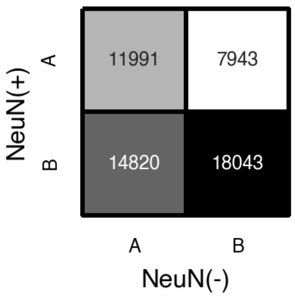


**Supplementary Figure S7.** Contingency table representing the distribution of 100-kb genomic bins between compartment types in NeuN(+) and NeuN(-) data.

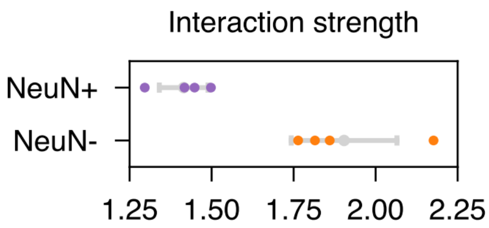


**Supplementary Figure S8**. Per-replicate compartment interaction strength calculated as the ratio of the highest intra-compartment interactions to the lowest inter-compartment interactions. Gray plots represent the mean and standard deviation of interaction strength across replicates.

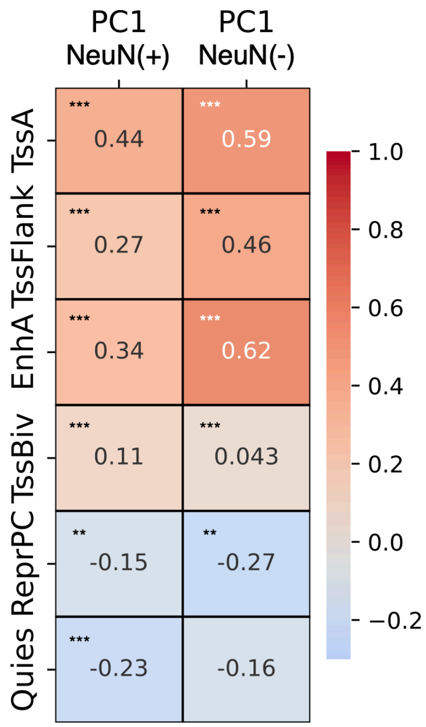


**Supplementary Figure S9.** Spearman's rank correlation coefficients between compartment eigenvectors (PC1) and chromatin chromHMM states retrieved from (4). Asterisks indicate correlation test p-values: *** – $p < 10^{-100}$, ** – $p < 10^{-20}$, * – $p < 10^{-4}$. Numbers indicate the correlation coefficients. Chromatin states annotation includes six distinct chromatin states, specifically EnhA (active enhancer), TssA (active promoter), TssBiv (bivalent promoter), TssFlnk (promoter flanking region), ReprPC (polycomb repression region), and Quies (other regions).

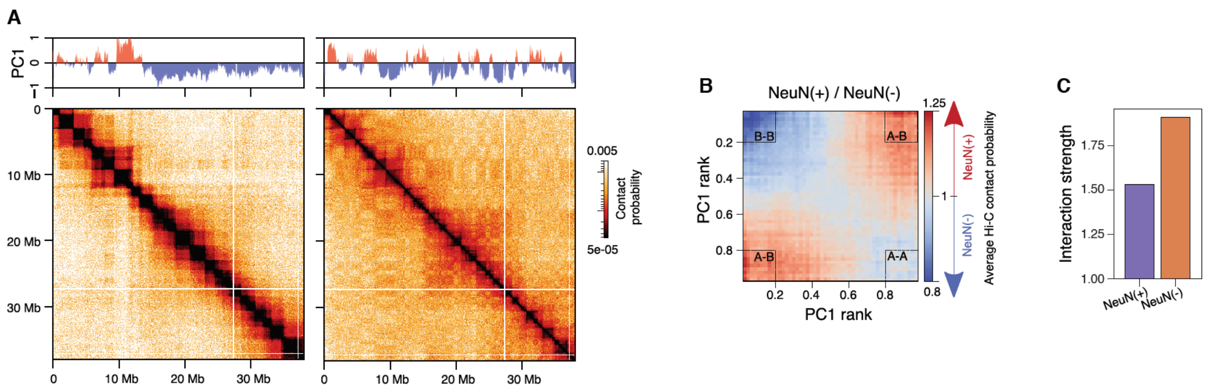


**Supplementary Figure S10**. Chromatin compartmentalization in cells derived from the prefrontal cortex (based on Hi-C data from Hu et al., 2021). (A) A fragment of the Hi-C map featuring compartment eigenvector (PC1) for NeuN(+) (left) and NeuN(-) (right) cells. Positive PC1 values represent the A compartment, while negative values correspond to the B compartment. (B) NeuN(+) / NeuN(-) ratio of average Hi-C contact probability (observed over expected) of genomic regions arranged by their respective PC1 rank (saddle plot). Both saddle plots were generated using NeuN(-) PC1. (C) Compartment interaction strength calculated as the ratio of the highest intra-compartment interactions to the lowest inter-compartment interactions.

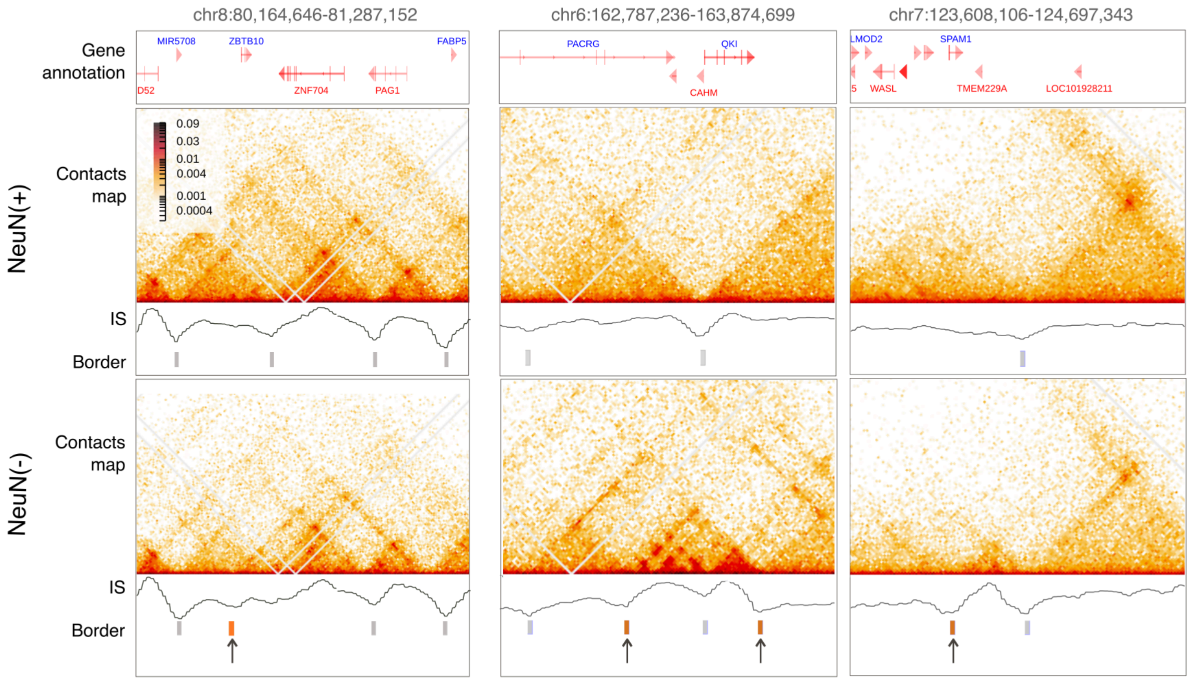


**Supplementary Figure S11**. Regions exhibiting differential NeuN(-) TAD borders. Contact maps for NeuN(+) and NeuN(-) cells are shown with the corresponding gene annotation, TADs borders layout, and insulation score track (IS).

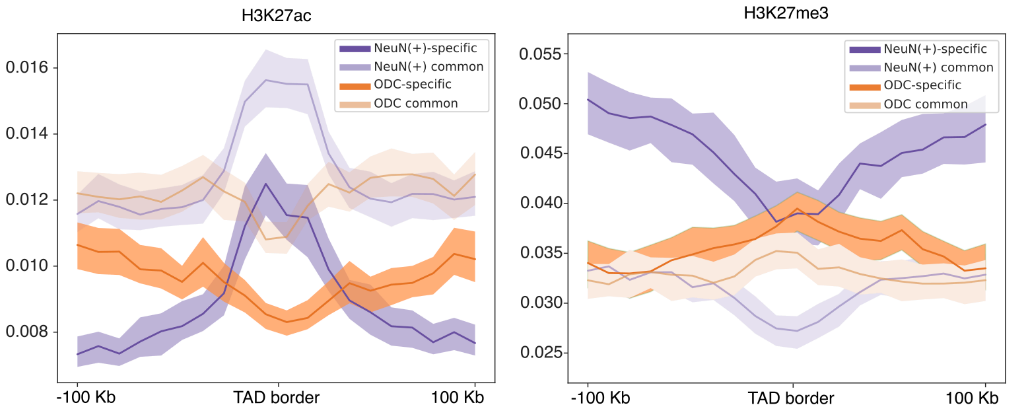


**Supplementary Figure S12**. H3K27ac and H3K27me3 histone mark profiles around the cell-type-specific and common TAD borders in oligodendrocytes.

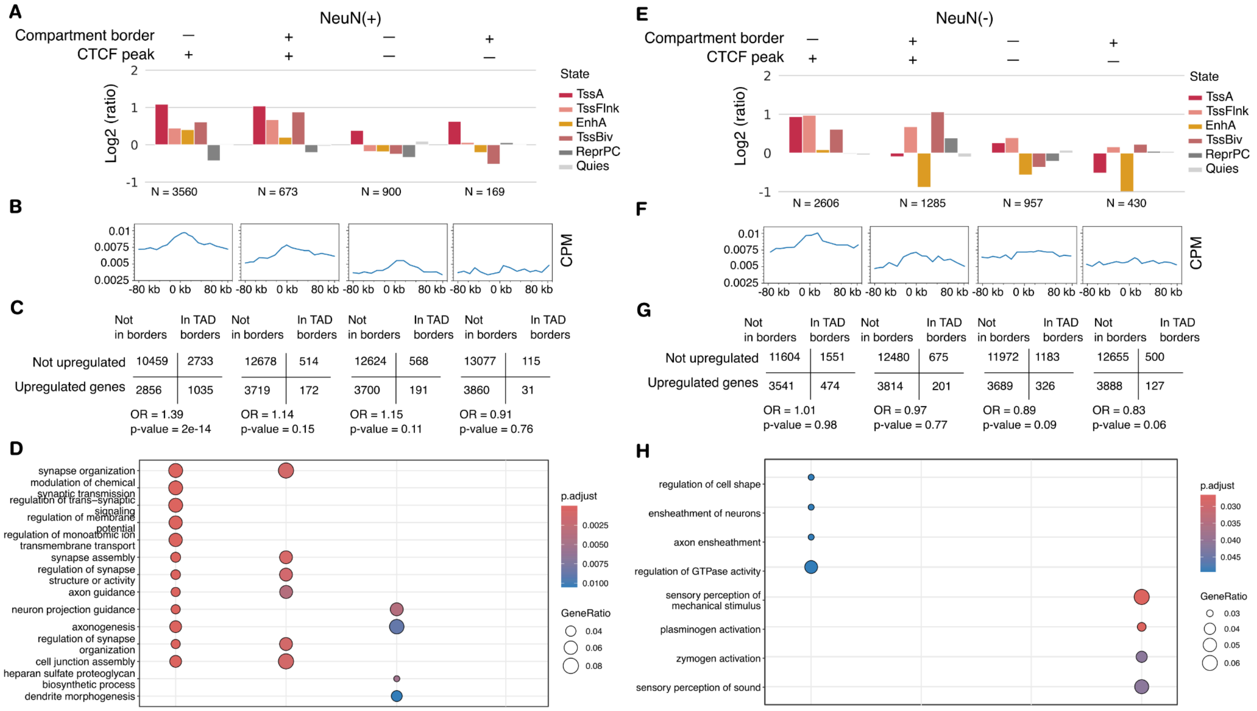


**Supplementary Figure S13**. Characteristics of TAD borders for NeuN(+) and NeuN(-) cells. Each of the figures A-H contains four columns corresponding to the TAD borders of different types. These TAD borders are categorized based on their association with CTCF peaks and compartment borders as follows: the first column includes TAD borders that coincide with CTCF peaks but not with compartment borders; the second column includes TAD borders that coincide with both CTCF peaks and compartment borders; the third column includes TAD borders that do not coincide with either CTCF peaks or compartment borders; and the fourth column includes TAD borders that coincide with compartment borders but not with CTCF peaks. **(A,E)** Grouped stacked bar plots of the chromatin state coverage within TAD borders for NeuN(+) (A) and NeuN(-) (E) cells. The ratio of each state is normalized to the total coverage within the genome in the corresponding cell type. Chromatin states annotation includes six distinct chromatin states, specifically EnhA (active enhancer), TssA (active promoters), TssBiv (bivalent promoters), TssFlnk (promoter flanking region), ReprPC (Polycomb repression region), and Quies (other regions). **(B,F)** Expression profiles around the cell-type-specific and common borders for NeuN(+) (B) and NeuN(-) (F) cells. **(C,G)** Confusion tables with Fisher test p-values for the enrichment of genes upregulated in NeuN(+) (C) or NeuN(-) (G) cells within corresponding TAD borders. DE genes were determined as genes upregulated in the selected cell type with FC > 1.5. **(D,H)** GO terms enrichment (shown with dots sizes) among upregulated in NeuN(+) (D) or NeuN(-) (H) genes located in neuronal-specific or common TAD borders.

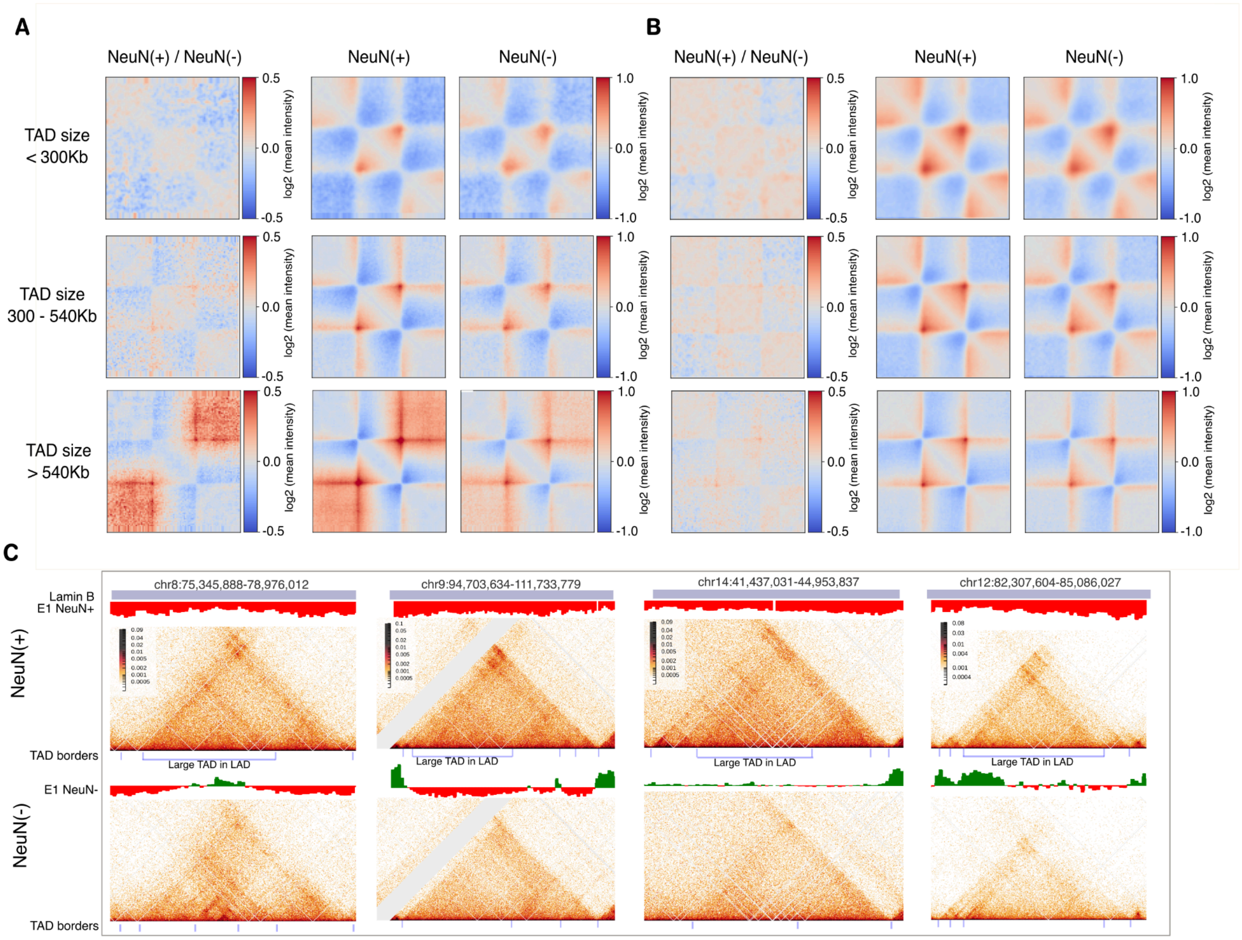


**Supplementary Figure S14.** Average TADs grouped by size. **(A)** TADs located within LADs and **(B)** outside of LADs. **(C)** The composition of large TADs within LADs. The track of Lamin B, E1 component, and the layout of TAD borders are shown for both NeuN(+) and NeuN(-) cell types.

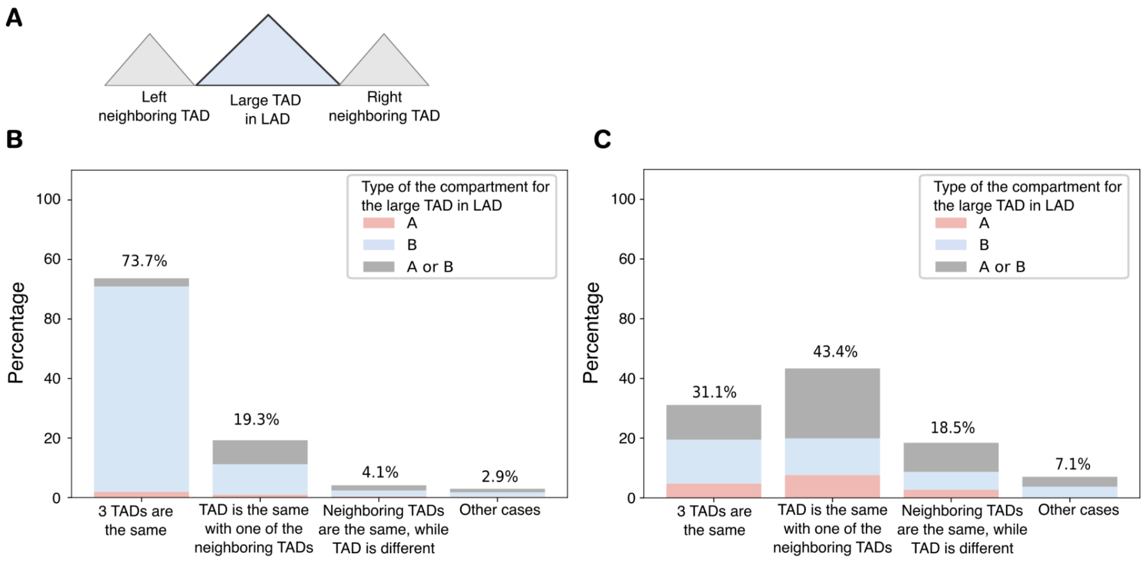


**Supplementary Figure S15**. Relationships between compartments and large TADs within LADs. (A) Schematic representation of the methodology used to analyze large TADs within LADs. (B) Distribution of TAD types categorized by compartment annotation in NeuN(+) cells. (C) Distribution of TAD types categorized by compartment annotation in NeuN(-) cells.

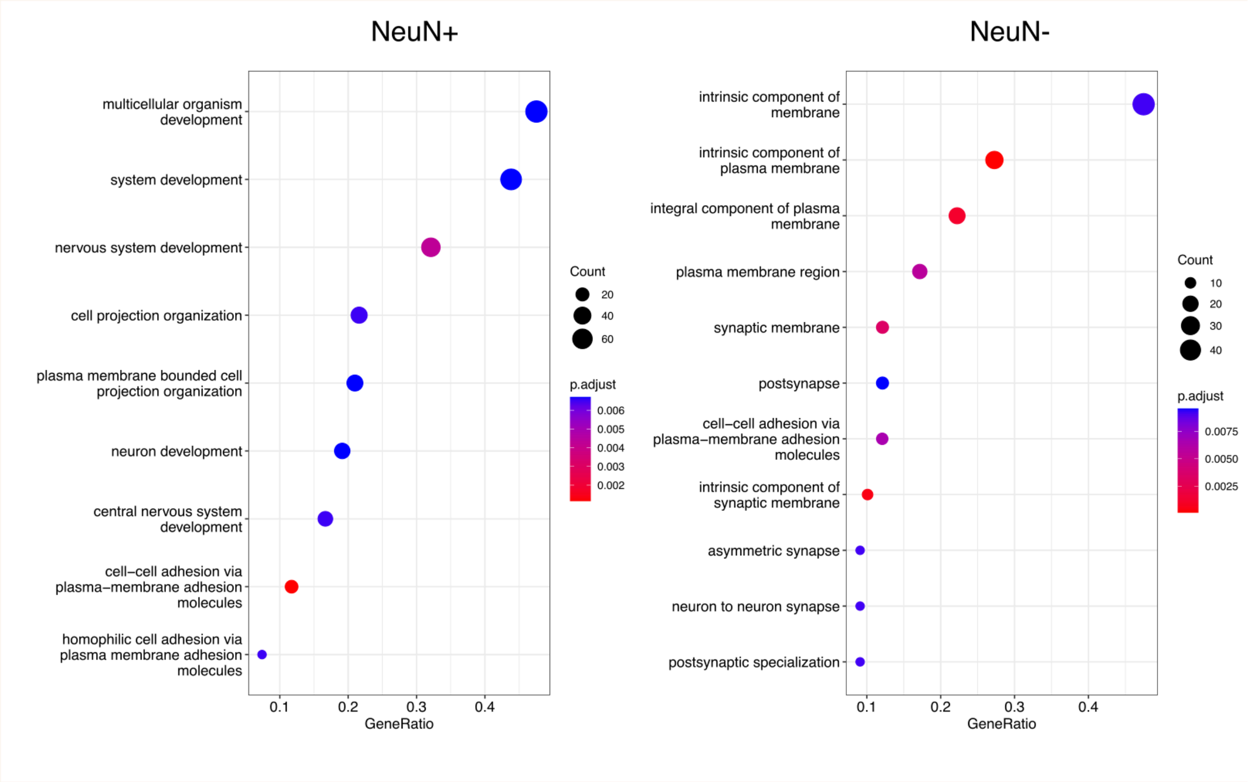


**Supplementary Figure S16.** GO terms enrichment (shown with dots sizes) among genes located at the borders of large TADs residing within LADs.

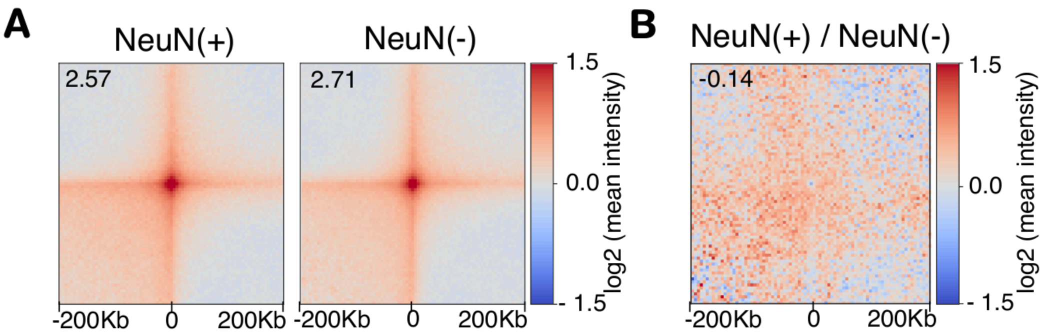


**C**

| NeuN(+) | Log2FC > 0.58 | Log2FC < -0.58 |
|---|---|---|
| In the loop | 294 | 217 |
| Not in the loop | 3597 | 3798 |

p-val = 1.2e-4

| NeuN(-) | Log2FC > 0.58 | Log2FC < -0.58 |
|---|---|---|
| In the loop | 216 | 367 |
| Not in the loop | 3675 | 3648 |

p-val = 1.3e-9

**Supplementary Figure S17.** Loops in NeuN(+) and NeuN(-) cells. **(A)** Average loop for NeuN(+) and NeuN(-) cells. **(B)** The ratio of average loops. **(C)** Contingency tables and Fisher's exact test p-values for the enrichment of loops with DE genes. Positive log2 fold change (Log2FC) corresponds to higher expression in NeuN(+).

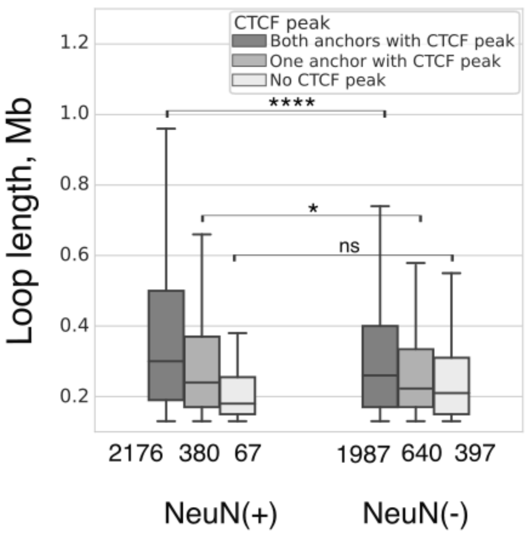


**Supplementary Figure S18**. Length of chromatin loops in NeuN(+) and NeuN(-) cells. Box plots illustrating the distribution loop lengths. Asterisks indicate Wilcoxon test p-values: **** - p < 0.00001.

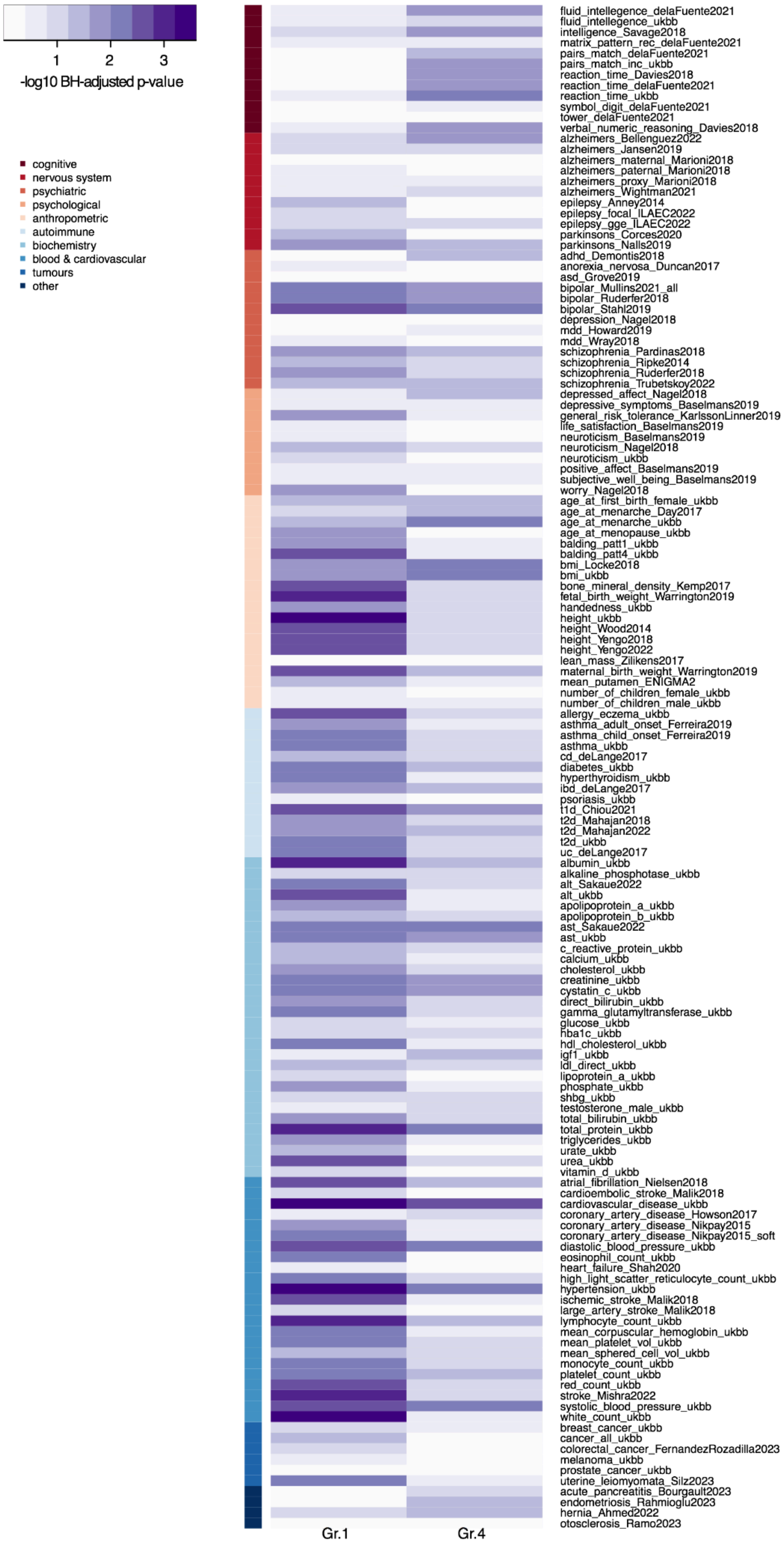
1
2
3
-log10 BH-adjusted p-value
cognitive
nervous system
psychiatric
psychological
anthropometric
autoimmune
biochemistry
blood & cardiovascular
tumours
other
fluid_intellegence_delaFuente2021
fluid_intellegence_ukbb
intelligence_Savage2018
matrix_pattern_rec_delaFuente2021
pairs_match_delaFuente2021
pairs_match_inc_ukbb
reaction_time_Davies2018
reaction_time_delaFuente2021
reaction_time_ukbb
symbol_digit_delaFuente2021
tower_delaFuente2021
verbal_numeric_reasoning_Davies2018
alzheimers_Bellenguez2022
alzheimers_Jansen2019
alzheimers_maternal_Marioni2018
alzheimers_paternal_Marioni2018
alzheimers_proxy_Marioni2018
alzheimers_Wightman2021
epilepsy_Anney2014
epilepsy_focal_ILAEC2022
epilepsy_gge_ILAEC2022
parkinsons_Corces2020
parkinsons_Nalls2019
adhd_Demontis2018
anorexia_nervosa_Duncan2017
asd_Grove2019
bipolar_Mullins2021_all
bipolar_Ruderfer2018
bipolar_Stahl2019
depression_Nagel2018
mdd_Howard2019
mdd_Wray2018
schizophrenia_Pardinas2018
schizophrenia_Ripke2014
schizophrenia_Ruderfer2018
schizophrenia_Trubetskoy2022
depressed_affect_Nagel2018
depressive_symptoms_Baselmans2019
general_risk_tolerance_KarlssonLinner2019
life_satisfaction_Baselmans2019
neuroticism_Baselmans2019
neuroticism_Nagel2018
neuroticism_ukbb
positive_affect_Baselmans2019
subjective_well_being_Baselmans2019
worry_Nagel2018
age_at_first_birth_female_ukbb
age_at_menarche_Day2017
age_at_menarche_ukbb
age_at_menopause_ukbb
balding_patt1_ukbb
balding_patt4_ukbb
bmi_Locke2018
bmi_ukbb
bone_mineral_density_Kemp2017
fetal_birth_weight_Warrington2019
handedness_ukbb
height_ukbb
height_Wood2014
height_Yengo2018
height_Yengo2022
lean_mass_Zilikens2017
maternal_birth_weight_Warrington2019
mean_putamen_ENIGMA2
number_of_children_female_ukbb
number_of_children_male_ukbb
allergy_eczema_ukbb
asthma_adult_onset_Ferreira2019
asthma_child_onset_Ferreira2019
asthma_ukbb
cd_deLange2017
diabetes_ukbb
hyperthyroidism_ukbb
ibd_deLange2017
psoriasis_ukbb
t1d_Chiou2021
t2d_Mahajan2018
t2d_Mahajan2022
t2d_ukbb
uc_deLange2017
albumin_ukbb
alkaline_phosphotase_ukbb
alt_Sakaue2022
alt_ukbb
apolipoprotein_a_ukbb
apolipoprotein_b_ukbb
ast_Sakaue2022
ast_ukbb
c_reactive_protein_ukbb
calcium_ukbb
cholesterol_ukbb
creatinine_ukbb
cystatin_c_ukbb
direct_bilirubin_ukbb
gamma_glutamyltransferase_ukbb
glucose_ukbb
hba1c_ukbb
hdl_cholesterol_ukbb
igf1_ukbb
ldl_direct_ukbb
lipoprotein_a_ukbb
phosphate_ukbb
shbg_ukbb
testosterone_male_ukbb
total_bilirubin_ukbb
total_protein_ukbb
triglycerides_ukbb
urate_ukbb
urea_ukbb
vitamin_d_ukbb
atrial_fibrillation_Nielsen2018
cardioembolic_stroke_Malik2018
cardiovascular_disease_ukbb
coronary_artery_disease_Howson2017
coronary_artery_disease_Nikpay2015
coronary_artery_disease_Nikpay2015_soft
diastolic_blood_pressure_ukbb
eosinophil_count_ukbb
heart_failure_Shah2020
high_light_scatter_reticulocyte_count_ukbb
hypertension_ukbb
ischemic_stroke_Malik2018
large_artery_stroke_Malik2018
lymphocyte_count_ukbb
mean_corpuscular_hemoglobin_ukbb
mean_platelet_vol_ukbb
mean_sphered_cell_vol_ukbb
monocyte_count_ukbb
platelet_count_ukbb
red_count_ukbb
stroke_Mishra2022
systolic_blood_pressure_ukbb
white_count_ukbb
breast_cancer_ukbb
cancer_all_ukbb
colorectal_cancer_FernandezRozadilla2023
melanoma_ukbb
prostate_cancer_ukbb
uterine_leiomyomata_Silz2023
acute_pancreatitis_Bourgault2023
endometriosis_Rahmioglu2023
hernia_Ahmed2022
otosclerosis_Ramo2023
Gr.1
Gr.4

**Supplementary Figure S19.** Heatmap of LDSC results for NeuN(-)-specific loops (Gr.1) and NeuN(+)-specific loops (Gr.4). Heatmap color is LDSC enrichment (–log10 p-values, p-values are BH adjusted). Rows are traits grouped in ten categories: cognitive, nervous system, psychiatric, psychological, anthropometric, autoimmune, biochemistry, blood & cardiovascular, tumours and other.

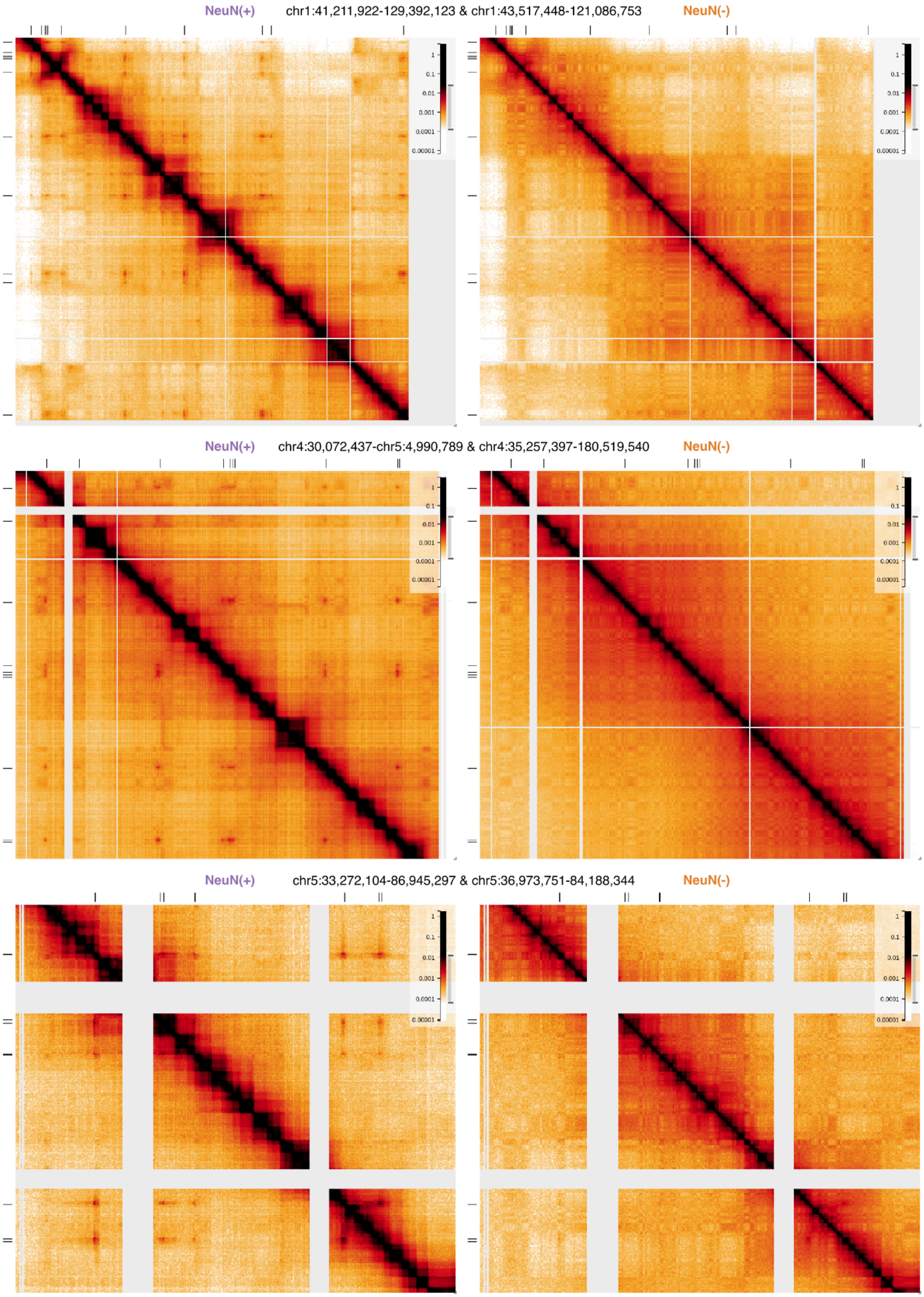


(see legend on the next page)

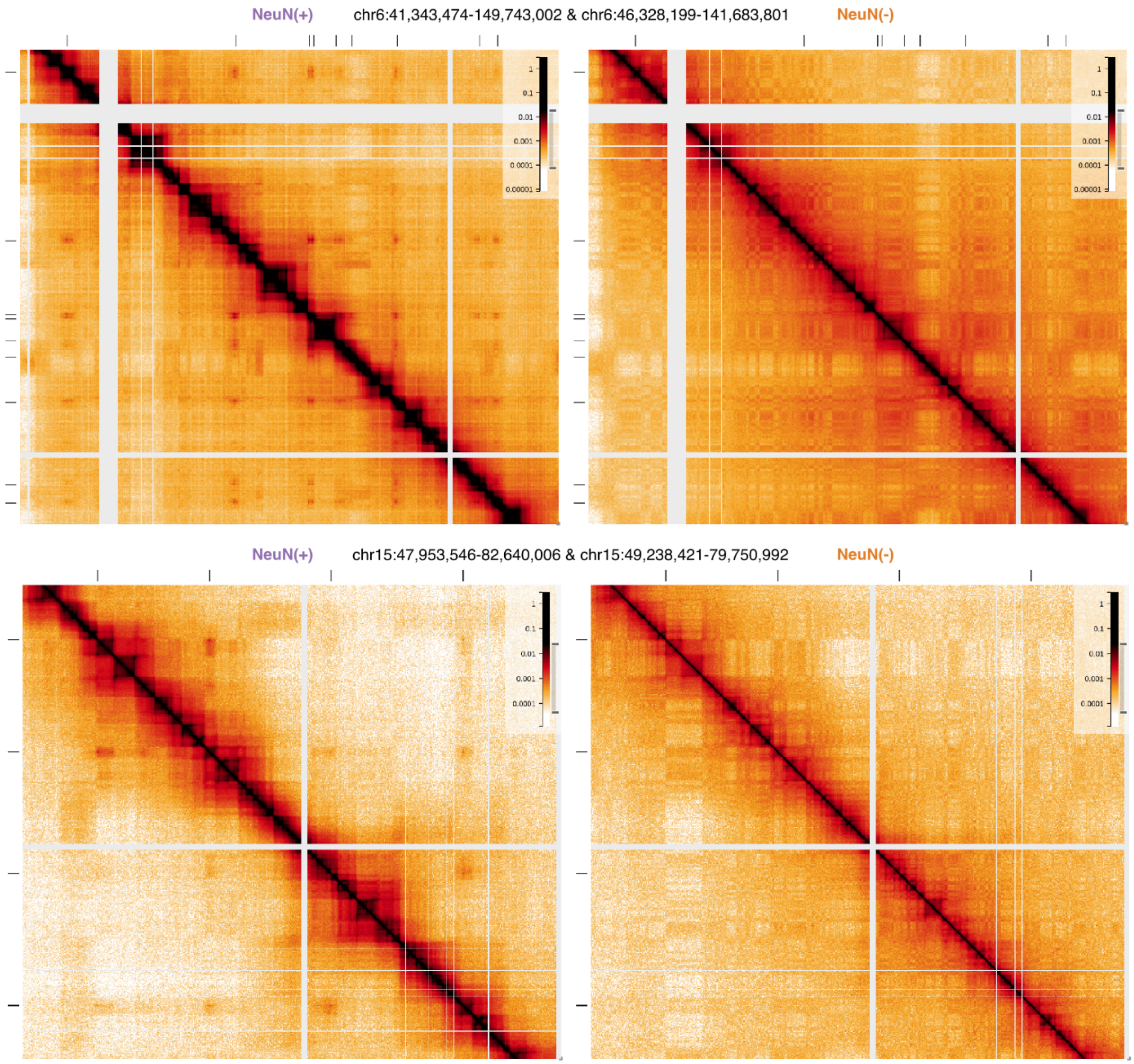


**Supplementary Figure S20.** Hi-C maps with examples of neuronal dots in NeuN(+) (left) and absence of dots in corresponding NeuN(-) (right) data. Track with black rectangles shows neuronal dot regions. Here Hi-C maps were merged with publicly available (Hu et al. 2021).

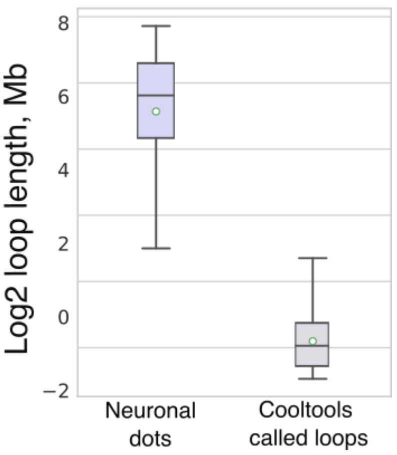


**Supplementary Figure S21**. Box plots of length distributions for neuronal dots (only cis-interactions are considered) and chromatin loops called using cooltools (NeuN(+) and NeuN(-) loops are combined).

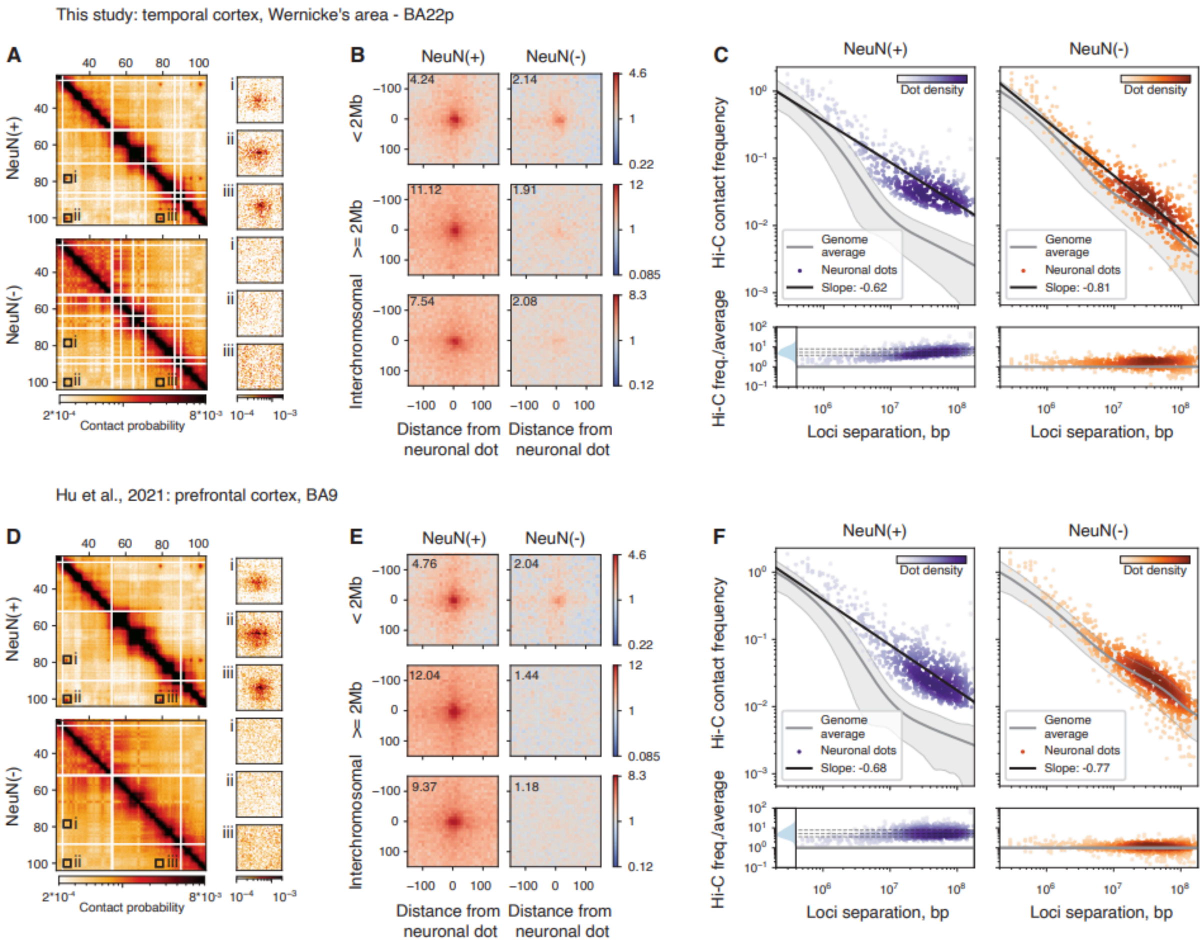


**Supplementary Figure S22**. Characteristics of neuronal dots in the temporal and prefrontal cortex areas. **(A,D)** A fragment of the Hi-C map with neuronal dots present in NeuN(+) (top), but absent in NeuN(-) (bottom) cells derived from the temporal (A) and prefrontal (D) cortex areas. **(B,E)** The average Hi-C signal (observed over expected) of neuronal dots (left) and the corresponding pairs of neuronal dot loci in NeuN(-) (right) cells derived from the temporal (B) and prefrontal (E) cortex areas. The value in the corner corresponds to the central pixel. **(C,F)** The contact scaling plots for cells derived from the temporal (C) and prefrontal (F) cortex areas. Left: the contact scaling of neuronal dots (purple circles), approximated by a linear regression (black line). Right: the contact scaling of pairs of neuronal dot loci in NeuN(-) (orange circles), also approximated by a linear regression (black line). The gray line represents the contact scaling of the whole genome.

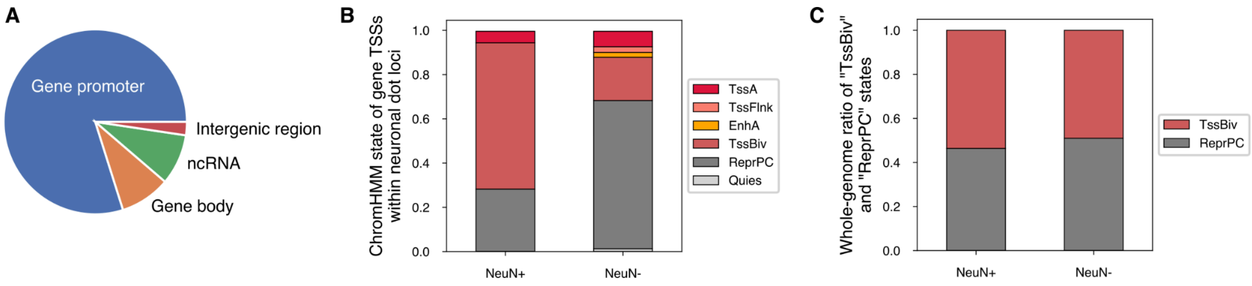


**Supplementary Figure S23.** Genes within neuronal dot loci. **(A)** Annotation of neuronal dot loci. **(B)** ChromHMM states of genes located at neuronal dot loci. **(C)** Whole-genome ratio of two ChromHMM states: “TssBiv” – “bivalent TSS”, occupied by H3K27me3 together with H3K4me3, and “ReprPC”, occupied by H3K27me3 without H3K4me3.

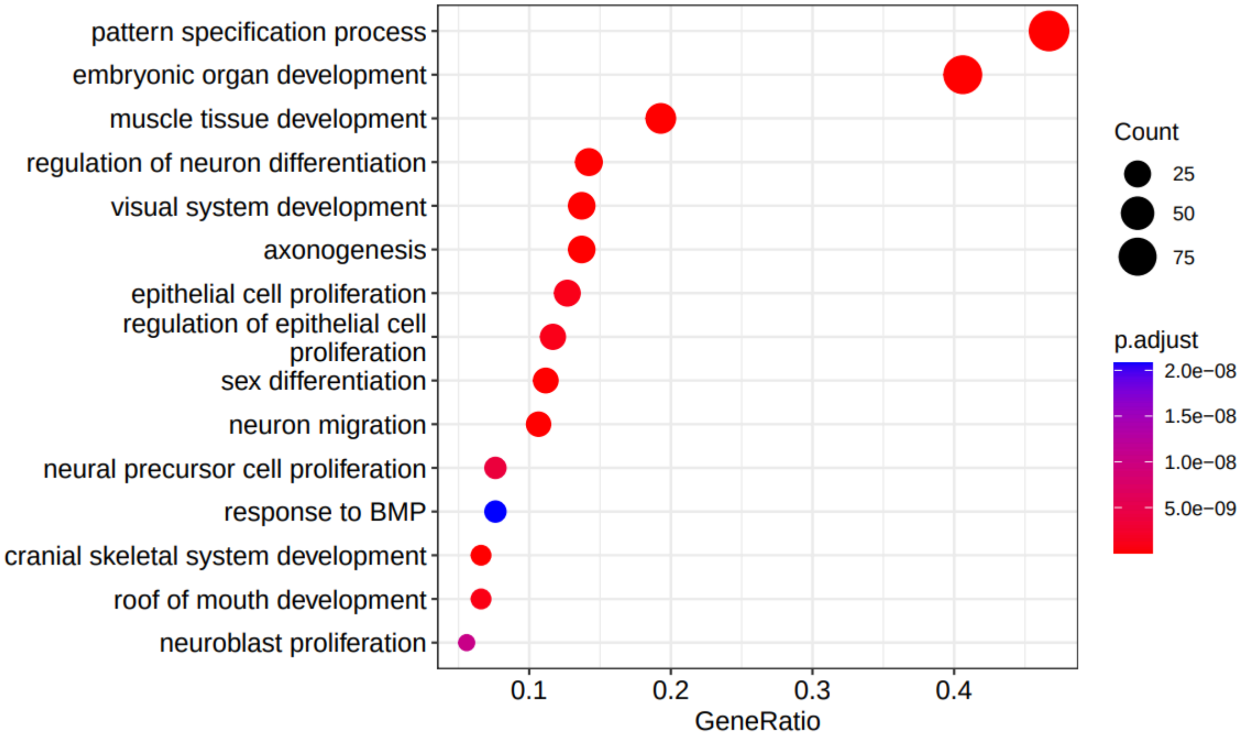


**Supplementary Figure S24.** Gene Ontology analysis of biological functions enriched in dot TFs. GeneRatio = k/n, where k – number of dot TFs falling in the particular GO category, n – all dot TFs.

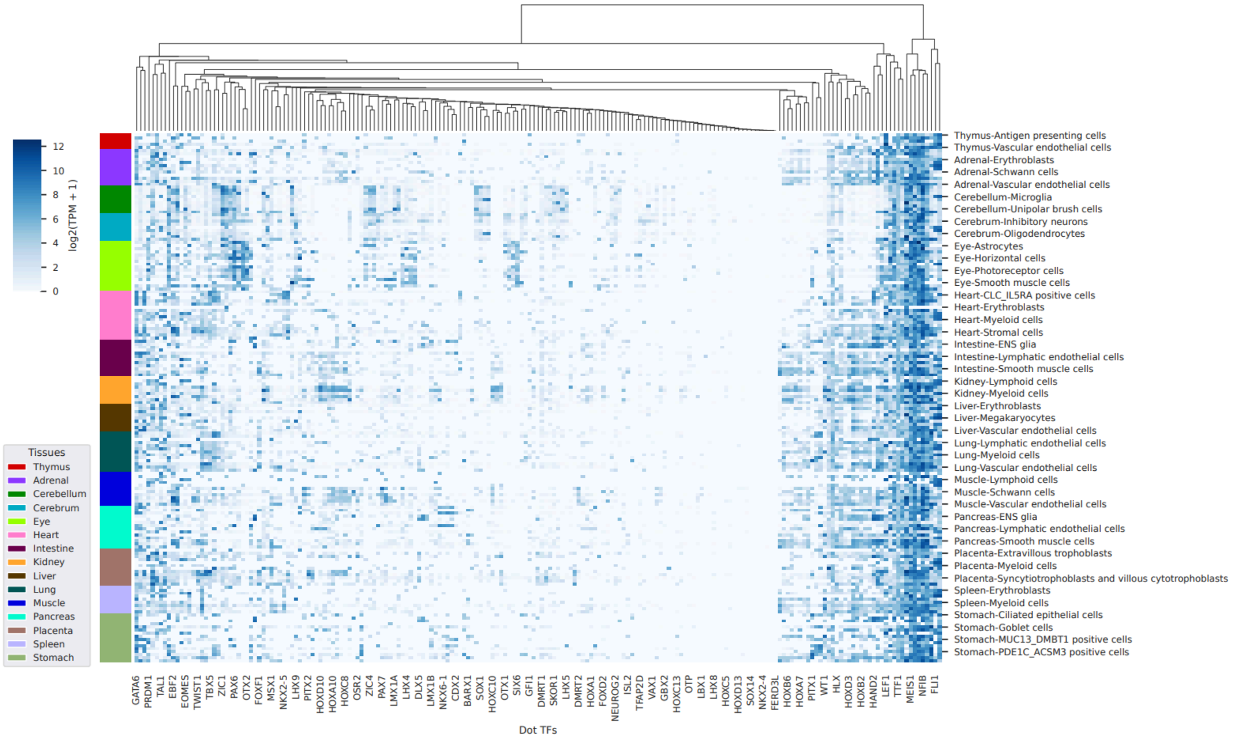


**Supplementary Figure S25.** Fetal expression of dot TFs in different tissues. scRNA-seq data obtained from (5).

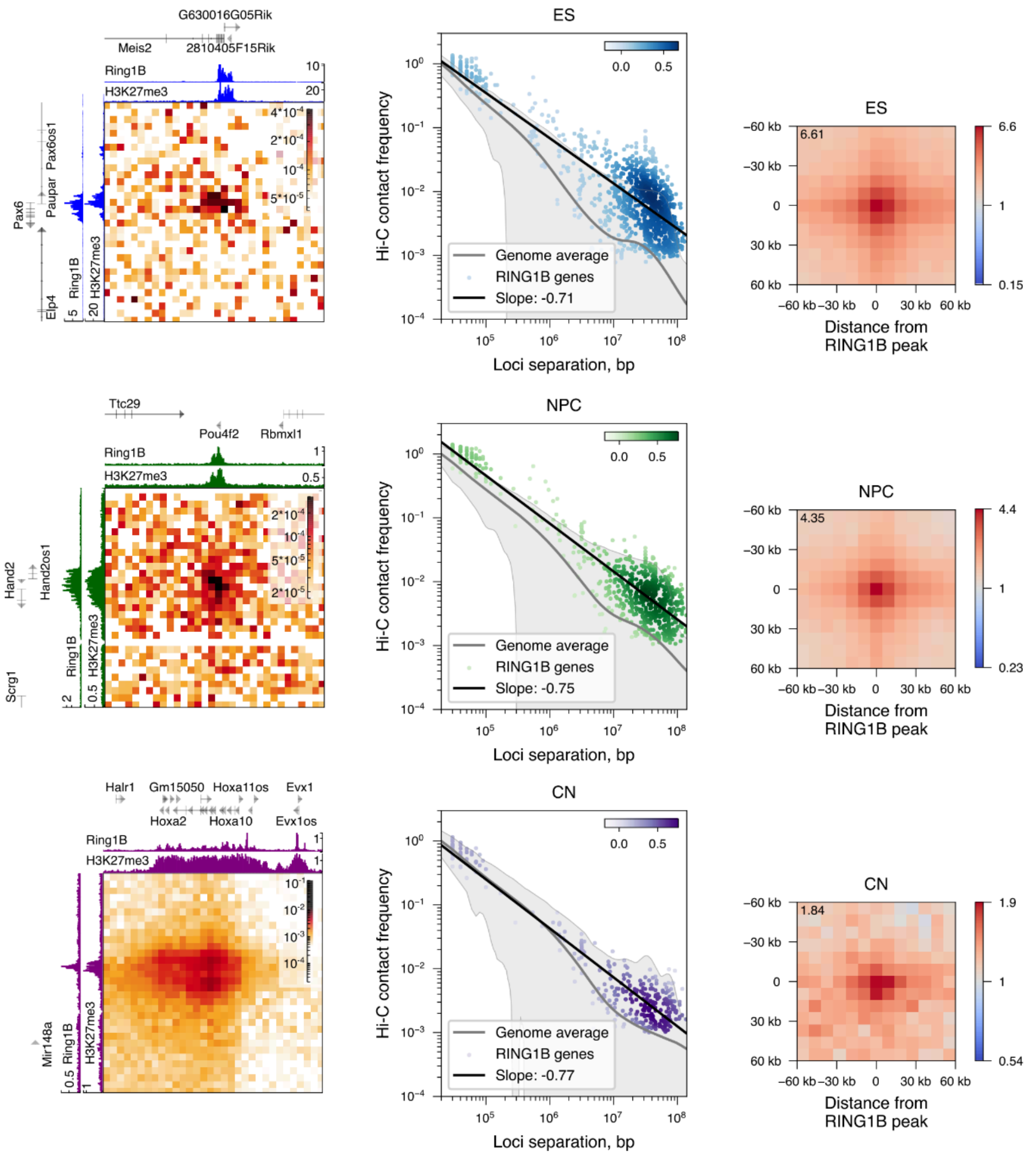


**Supplementary Figure S26.** Mouse orthologs of neuronal dot genes form long-range, RING1B-coupled interactions in embryonic stem cells (top row), neural progenitor cells (middle row) and cortical neurons (bottom row). Left column: a region of a Hi-C map showing interaction between mouse orthologs of neuronal dot genes, with corresponding gene annotation and ChIP-seq tracks of H3K27me3 and RING1B. Center column: contact scaling of mouse orthologs of neuronal dot genes overlapping RING1B ChIP-seq peaks, approximated by a linear regression – black line. Grey line – whole-genome expected. Right column: average Hi-C signal of pairs of mouse orthologs of neuronal dot genes overlapping RING1B ChIP-seq peaks (observed over expected). Value in the corner corresponds to the central pixel. Reanalysed data from (6).

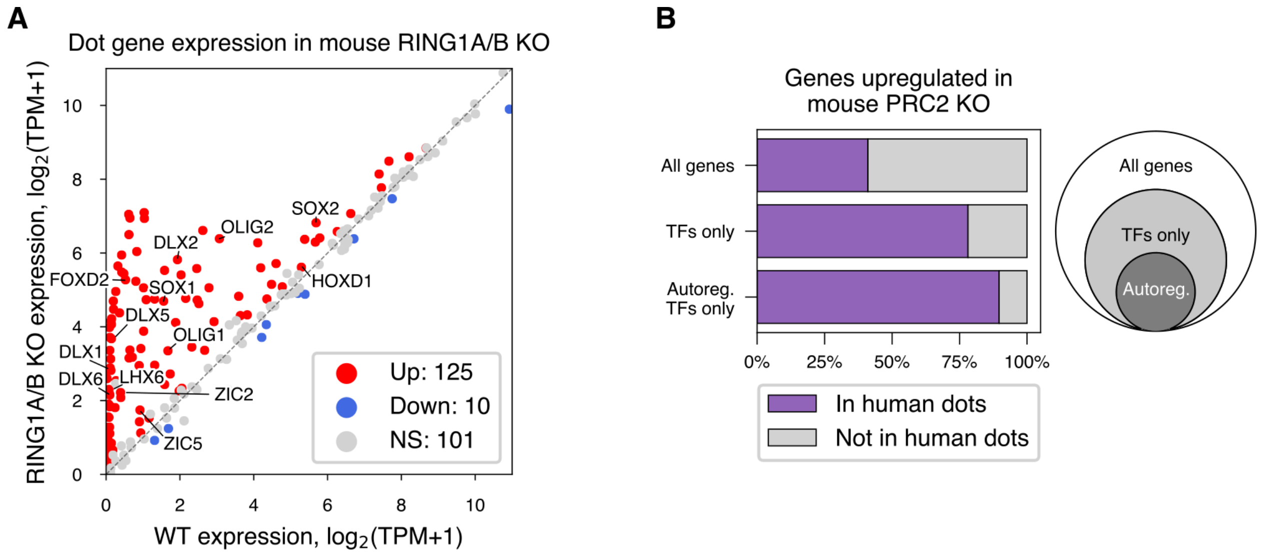


**Supplementary Figure S27.** Impact of polycomb knockouts on neuronal dot gene expression. **(A)** Expression of mouse orthologs of neuronal dot genes in wild-type (WT) and RING1A/B knockout (KO) of mouse motor neurons. Data from Sawai et al.(7). Colors represent upregulated (Up), downregulated (Down) and not differentially expressed (NS) groups of genes with corresponding group sizes. 9 out of 245 dot gene orthologs were not present in Sawai et al. data, thus, the total number of genes here is 236. **(B)** Overlap between human dot genes and their mouse orthologs that are upregulated in mouse PRC2 knockout. Data from Von Schimmelmann et al.(8). Overlap is shown for three sets of upregulated genes: all genes, TFs only and TFs with positive auto-regulation only.

**Supplementary Table S1.** Sample metadata.

| Cell type | Sample name | Sex | Age, years | Total number of reads | Number of contact pairs after filtering |
|---|---|---|---|---|---|
| NeuN+ | Sample_1 | female | 59 | 235,775,789 | 132,609,155 |
| NeuN- | | | | 224,307,550 | 103,174,134 |
| NeuN+ | Sample_2 | male | 58 | 181,179,355 | 102,847,525 |
| NeuN- | | | | 248,076,415 | 78,650,805 |
| NeuN+ | Sample_3 | female | 62 | 200,429,327 | 104,608,807 |
| NeuN- | | | | 178,676,161 | 96,022,439 |
| NeuN+ | Sample_4 | female | 36 | 186,290,474 | 85,560,365 |
| NeuN- | | | | 219,229,754 | 109,353,761 |
